# First Evidence of Dicistroviruses Infecting Protists

**DOI:** 10.64898/2026.07.25.740690

**Authors:** Julie Thomy, Christopher R. Schvarcz, Kelsey A. Allen, Kyle F. Edwards, Grieg F. Steward

## Abstract

Molecular surveys suggest that RNA viruses are abundant and diverse in the ocean, but the hosts for most of these viruses are unknown because so few have been cultivated. Here, we present the genomes of five positive-sense, single-stranded RNA (+ssRNA) viruses isolated from tropical seawater that infect green algae in the genus *Tetraselmis* (order Chlorodendrales). Phylogenetic analyses of multiple genes placed these closely related viruses within the family *Dicistroviridae* (order *Picornavirales*) making them the first viruses within the bounds of the *Dicistroviridae* family demonstrated to infect an organism other than arthropods. The RNA-dependent and capsid gene sequences of the *Tetraselmis* RNA viruses (TetRNAV01–05) cluster with others recovered from diverse environmental water samples or aquatic invertebrate tissues, and together they form a strongly supported sister clade to viruses in the genus *Triatovirus*. The TetRNAVs harbor a unique intergenic internal ribosome entry site (IRES), suggesting a translation strategy distinct from that described in arthropod-infecting dicistroviruses. Our results suggest that many uncultivated viruses presumed to infect invertebrates, because they were detected in invertebrate-derived samples and their genes cluster within the family *Dicistroviridae*, may instead be protist-infecting viruses.

## Introduction

Marine viruses strongly influence microbial mortality, food web dynamics, and biogeochemical cycling of carbon and nutrients (1). Much of the early research on viruses infecting eukaryotic algae focused on viruses with double-stranded DNA genomes (2–4) but the prevalence and importance of marine RNA viruses is becoming better appreciated. In 2003, the isolation of a positive-sense, single-stranded RNA ((+)ssRNA) virus infecting the raphidophyte *Heterosigma akashiwo* provided the first clear evidence of an RNA virus infecting a marine protist (5, 6). Subsequently, numerous ssRNA viruses, infecting a variety of protists including dinoflagellates, thraustochytrids, and diatoms, have been isolated from a broad range of marine environments (7–17). Most of these are members of the family *Marnaviridae* within the order *Picornavirales*. A smaller portion of protist virus isolates are grouped within the families *Alvernaviridae* (the order *Sobelivirales*) and *Reoviridae* (the order *Reovirales*) (8, 18). Rapid advances in metagenomic and metatranscriptomic approaches have revealed an extraordinary diversity of RNA viruses in aquatic ecosystems across temperate (19–21), tropical (22–25), and polar regions (26–29). Two studies have further suggested that aquatic RNA viruses are not only diverse, but, at least in coastal tropical and polar waters, their abundance was comparable to that of DNA viruses (26, 30). A common theme of the numerous surveys of aquatic RNA virus diversity is that (+)ssRNA viruses classified within the order *Picornavirales* dominate the aquatic RNA virosphere.

The order *Picornavirales* comprises small, non-enveloped viruses characterized by icosahedral capsids approximately 30 nm in diameter that infect a wide range of eukaryotic hosts, including vertebrates (animals and humans), invertebrates (arthropods and insects), as well as protists and plants (31). Reflecting this broad host range, *Picornavirales* is recognized as a viral supergroup that, according to the International Committee on the Taxonomy of Viruses (ICTV, 2019), encompasses the families *Picornaviridae, Dicistroviridae, Iflaviridae, Marnaviridae, Secoviridae, Caliciviridae, Solinviviridae, Noraviridae,* and *Polycipiviridae*. Members of the order *Picornavirales* (i.e., picornavirads) share a common genomic architecture based on a (+)ssRNA genome, typically ranging from 7 to 12.5 kilobases in length that serves directly as a template for protein synthesis. Like cellular mRNAs, their genomes are generally flanked by a long 5′ untranslated region (UTR) and a 3′ polyadenylated (poly(A)) tail. However, picornavirad genomic RNA lacks a 5′ cap structure and instead has a small viral protein (VPg) covalently attached to the 5′ end (31), highlighting a key molecular adaptation for host exploitation.

The shared genomic features of picornavirads reflect their common origin, but extensive diversification driven by recombination, horizontal gene transfer, host adaptation, and genome rearrangements has resulted in the emergence of multiple distinct lineages (32–35). Picornavirad genomes can be organized either as a single open reading frame (monocistronic), encoding one large polyprotein, or as two distinct open reading frames (dicistronic), each producing separate polyproteins. These genomes encode both non-structural replication proteins—such as the RNA helicase (Hel), cysteine protease (Pro), and RNA-dependent RNA polymerase (RdRP)—and structural capsid proteins (VP1, VP2, VP3, and VP4), which follow distinct evolutionary trajectories (31, 33). Another distinct feature of these viruses is the presence of long, highly structured untranslated regions that function as internal ribosome entry sites (IRES). These elements enable cap-independent initiation of translation, allowing the virus to hijack host translational machinery. In most *Picornavirales*, the IRES is located within the 5′ UTR of a single ORF. In contrast, members of the *Dicistroviridae* family (dicistrovirids) have evolved a unique strategy, with an additional IRES located in the intergenic region (IGR) (36–39). This dual-IRES system enables independent regulation of non-structural and structural protein synthesis during infection, with replication enzymes typically produced early and structural proteins synthesized at later stages.

Despite the large number of environmental studies on aquatic RNA viruses, significant gaps remain in our understanding of their diversity, distribution, replication strategies, and host interactions, particularly for those infecting protists. Characterization of new isolates can be particularly helpful for understanding ecological and evolutionary questions. Here, we report the first evidence of small (approximately 35 nm in diameter) ssRNA viruses infecting marine algae in the genus *Tetraselmis*. The algae belong to the family Chlorodendrophyceae within the phylum Chlorophyta (green algae) and are part of the core Chlorophyta lineage (40, 41). Members of the genus *Tetraselmis* are unicellular green algae (6–10 μm) characterized by a thin, scaly cell wall, four anterior flagella covered with mastigonemes, and a single posterior chloroplast containing both an eyespot and a pyrenoid. Species of *Tetraselmis* are widely distributed across marine, brackish, and freshwater environments (42–44). Although they are primarily free-living, some species occur as endosymbionts of acoel flatworms (*Symsagittifera roscoffensis*, originally known as *Convoluta roscoffensis*) or of radiolarians such as *Spongodrymus* (45–48). In addition, *Tetraselmis* species are extensively used in aquaculture as a food source for juvenile molluscs, shrimp larvae, and rotifers (49–51). While numerous RNA viruses have been isolated and or detected from the tissues of marine invertebrates such as arthropods and mollusks (mainly associated with the order *Picronavirales*), only DNA viruses have so far been reported to infect the green alga *Tetraselmis* (52–54). Our results identify the newly isolated viruses as members of the order *Picornavirales*, classified within the family *Dicistroviridae*—a group previously known only to infect animals in the phylum Arthropoda.

## Material and methods

### Isolation and maintenance of algae and viruses

Nine *Tetraselmis* spp. strains (class Chlorodendrophyceae) were isolated from tropical marine water collected on multiple occasions from 2010–2011 by enrichment of whole or filtered (< 5 µm) seawater with f/2 nutrients (55, 56) followed by three consecutive rounds of serial dilution to extinction to obtain unialgal cultures (52) (Table S1). Six strains (KB-FL30, KB-FL40, KB-FL45, KB-FL39, KB-CR05, KB-FL46) were isolated from coastal waters of Kāne‘ohe Bay (21°25’46.80”N, 157°47’31.51”W) and three (AL-FL03, AL-FL22 and AL-FL18) from open ocean waters of Station ALOHA (22°45’ N, 158°00’ W) in the North Pacific Subtropical Gyre. Strains from Station ALOHA were then maintained in K medium (57), and those from Kāneʻohe Bay in f/2 medium, by transferring inocula of dense culture (0.5 mL) into fresh medium (20 mL) in glass tubes every one to two months. Cultures were grown at 24°C, under a 12:12 light:dark cycle and with an irradiance of approximately 30–100 μmol photons m^−2^ s^−1^. The strains have been maintained as part of a boutique collection of marine protistan plankton at the University of Hawai’i at Mānoa (UHM) and were assigned collection identifiers (Table S1).

To isolate viruses infecting these strains, a large volume of seawater (200 liters) was collected from the same location and the same time as the green alga isolates described above. The water samples were pre-filtered (0.8 µm polycarbonate track-etched membrane, Isopore; Millipore) to remove larger cells and particles. Then, small particles were concentrated by tangential flow filtration (TFF; Pellicon 2 Mini System; Millipore) using filters with a nominal molecular weight limit (NMWL) of 30 kilodaltons (kDa). Concentrates were amended with either K or f/2 nutrients to match the medium of the alga being challenged, then added to the algal cultures at 1:20 dilution. Only virus concentrated from Kāne‘ohe Bay caused lysis of any of the *Tetraselmis* strains. After propagating the *Tetraselmis* lysates for multiple generations, the viruses were isolated using three rounds of dilution-to-extinction performed in 96-well plates (58). A total of nine virus isolates has since been maintained by serial transfer of lysate every one to two months into a culture of the host alga on which each virus was first isolated. After seven days, a portion of the lysed culture is archived at 4°C and used as the inoculum for the next challenge. One of the isolates, TetV-1, was described elsewhere (52).

### Virus concentration and purification

For subsequent transmission electron microscopy (TEM) and host range analyses, algal cultures in exponential growth phase (70 mL) were inoculated with viral lysate (2 mL). After five days, the clear lysates were filtered through 0.6 µm polycarbonate membrane and concentrated by centrifugal ultrafiltration (Vivaspin 100, 100 kDa; Sartorius) at 2,000 × g for 20 min. The concentrated lysate (approximately 300 µL) was purified on a continuous equilibrium buoyant density gradient prepared with iodixanol (OptiPrepTM, Serumwerk Bernburg AG) as previously described (59). Briefly, pre-formed gradients were made with low (20%) and high-density (40%) solutions prepared in 0.2 μm (Whatman Anotop, 25 mm) filtered, autoclaved filtered seawater. Concentrated lysates were layered on top of the pre-formed gradients in a 13.2 mL ultraclear tube (Beckman Coulter, ref: 344059) and centrifuged at 35,000 RPM (210,000 × *g*; SW 41 Ti rotor) at 20°C for approximately 20 hours. Each gradient fraction of 500 µL was collected top end first using the BioComp piston fractionator (Model 153). The density of fractions was determined by measuring the mass using a precision balance and a micropipette with disposable, positive-displacement tips (59). Iodixanol was buffer exchanged three times with 0.02 μm-filtered (Whatman Anotop, 25 mm), autoclaved seawater by repeated centrifugal ultrafiltration (Amicon Ultra-0.5 100 kDa NMWL; Millipore) at 14,000 × *g* for 10 min. Fractions ranging between 1.10 and 1.14 g/mL were checked for the presence of viral particles by TEM.

### Transmission Electron Microscopy

The morphology of viral particles was observed by electron microscopy after negative staining with uranyl acetate. Briefly, 200-mesh copper grids with carbon-stabilized formvar supports (25-50 nm Formvar and 3-4 nm Carbon) were glow discharged (Denton Vacuum DV-502A) for 2 minutes. Four microliters of purified viral samples were deposited onto the grid. After 45 sec, the sample was carefully wicked away with filter paper and four microliters of 2% uranyl acetate was applied to the grid for 45 seconds. The stain was carefully wicked away and the grid immediately washed by depositing 4 µL of distilled water, which was immediately wicked away. The grid was left to dry for a minimum of two hours before viewing on a Hitachi HT7700 TEM at 100 kV and photographed with an AMT XR-41B 2k x 2k551 CCD camera. The size of viral particles (n = 100) was measured using the tool ImageJ (v1.54) (60).

### Host range analysis

Host range was tested on eight *Tetraselmis* spp. strains. Each infectivity test was performed in triplicate in 96-well plates. For challenged wells, 200 µL of alga culture in late exponential phase was inoculated with 2 µL of fresh viral lysate either unfiltered or filtered through a 0.1 µm syringe filter (Acrodisc, Pall). Control wells were mock inoculated with 2 µL of medium (K or f/2) depending on the host tested. The algal cells were monitored for lysis daily for seven days by quantifying chlorophyll *a* autofluorescence in a microplate reader (PerkinElmer). Taxonomic classification of the tested microalgae was reported in a previous study based on 18S rRNA gene sequences in the NCBI database (52) (Table S1).

### Nucleic acids extraction and identification

Exponentially growing algal cultures (30 mL) were inoculated with 1 mL of lysate containing purified virions. After five days, the lysate was centrifuged at 6,000 × *g* for 15 min at 20°C to pellet cell debris and the supernatant was filtered through 0.45 µm (Syringe filter, cellulose acetate membrane, Avantor) or 0.1 µm (Syringe filter, PES, Acrodisc), for larger and small viruses, respectively, as determined by electron microscopy. Twelve milliliters of the filtrate was concentrated using centrifugal ultrafiltration (Amicon Ultra-4, 100 kDa NMWL,; Millipore) at 4,000 × *g* in a swinging bucket for 15 min at 20°C. Total nucleic acids were extracted from concentrated viral lysates (100–175 µL) using MasterPure™ Complete DNA & RNA Purification Kit (LGC Biosearch, ref: MC85200) following manufacturer’s instructions. Briefly, 1 volume of 2× T and C Lysis Solution mixed with 1 µL of proteinase K (50 μg μL^-1^) was added to the sample and incubated at 65°C for 30 min. After being placed on ice, a 1:2 volume of MPC Protein Precipitation Reagent was added to the lysed sample and mixed vigorously. The debris was then centrifuged at 10,000 × *g* for 10 min at room temperature. The supernatant was then collected and mixed with one volume of isopropanol. The mixture was incubated overnight at 4°C and the nucleic acids were pelleted by centrifugation at 15,000 × *g* for 30 min. The pellet was washed three times with 70% ethanol. After complete drying, the nucleic acids were resuspended in 35 µL of nuclease-free water.

For each virus, the extracted nucleic acids were subjected to enzymatic digestion to determine their composition. For DNA digestion, 1 µL of DNase I (1 U µL^-1^), 3 µL of 10× Reaction Buffer with MgCl₂, and 2 µL of sample were combined with nuclease-free water to a final volume of 30 µL. For RNA digestion 1 µL of RNase A (5 μg μL^-1^), 3 µL of 10× Reaction Buffer, and 2 µL of sample were combined with nuclease-free water to a final volume of 30 µL. Samples were purified on a column using DNA Clean & Concentrator-5 kit (Zymo Research) to remove enzymes and debris. Samples were denatured for 10 minutes at 70°C in 2× RNA loading dye (New England Biolabs) and then quick-chilled on ice. Denatured samples were loaded onto 1% agarose-formaldehyde gel prepared with 1× MOPS buffer (0.02 M 3-(N-morpholino) propanesulfonic acid, 2 mM sodium acetate, 1 mM EDTA, pH 7) and 2.2 M formaldehyde and run in 1× MOPS buffer at 65V for 75 min. The gel was stained in a bath of Diamond™ Nucleic Acid Dye (Promega) for 30 min and visualized on a blue-light transilluminator (Thermo Fisher Scientific). Of the nine virus isolates, five were determined to have RNA genomes and were assigned strain names of TetRNAV01, TetRNAV02, TetRNAV03, TetRNAV04 and TetRNAV05. Of the DNA-containing viruses, one was described previously (Schvarcz and Steward, 2018) and the other three will be presented elsewhere.

For genome sequencing, total RNA was extracted from TetRNAV02–05 lysates using the protocol described above. After treatment with DNase I (1 U µL^-1^), RNA was column purified (RNA Clean & Concentrator-5 kit; Zymo Research) and eluted in 30 µL of nuclease-free water. The concentration of purified total RNA extract was determined by fluorometry (Qubit RNA HS Assay Kit;Thermo Fisher Scientific). For TetRNAV01 virus, the process was similar, but a larger volume of lysate (20 L) was processed, and the virions were purified by banding in CsCl (61). TetRNV01 lysate was clarified by centrifugation at 4,000 × *g* for 30 min. The supernatant was concentrated using tangential flow filtration with a 30 kDa NMWL cassette filter; (Pellicon 2 Mini system; Millipore), then the concentrate was centrifuged a second time at 4,000 × *g* for 30 min and the supernatant filtered through a 0.22 μm pore size polyethersulfone membrane filter (Sterivex; Millipore). Finally, a last concentration was performed using a centrifugal ultrafilter (30 kDa or 100 kDa NMWL Centricon Plus-70; Millipore). Finally, viral particles were purified by banding in a cesium chloride (CsCl) equilibrium buoyant density gradient (59). After collecting the virus-purified band from the CsCl gradient, the sample was buffer exchanged three times with SM buffer (Sodium Chloride, Magnesium sulfate, Tris buffer) using centrifugal ultrafiltration (30 kDa NMWL filters; Millipore Amicon Ultra-0.5) and total nucleic acids were extracted from the concentrates as described above.

### Random-priming sequence-independent single-primer amplification

For TetVRNAV02–05, complementary DNA (cDNA) synthesis was performed by reverse transcription (LunaScript® RT Master Mix Kit, Primer-free; New England Biolabs) by combining one microliter of purified viral RNA (approximately 12 ng), and primer K-8N (5’-GACCATCTAGCGACCTCCACNNNNNNNN-3’) and nuclease-free water to a final volume of 14 µL. The mixture was heated to 65°C for 5 min to disrupt RNA secondary structure, then placed on ice for 1 min. LunaScript RT master mix was then added to the tube and mixed gently (20 µL final volume). The reaction was incubated at 25°C for 2 min to allow primer annealing, followed by cDNA synthesis at 55°C for 10 min and enzymatic inactivation at 95°C for 1 min. The second DNA strand was synthesized by adding 20 µL of the first-strand cDNA to 10 pmol of primer K-8N, and 400 μM dNTPs in 1× Klenow reaction buffer (New England Biolabs). Subsequently, 1 µL of Klenow fragments (200 U) (New England Biolabs) was added and incubated at 37°C for 60 min (final volume, 25 µL). For the sequence-independent PCR amplification, 5 µL of double-stranded cDNA (ds cDNA) was added to a reaction mix containing 1× Platinum ™ SuperFi™ II Green PCR Master buffer (Thermo Fisher Scientific) and 1 µM of primer K (5’-GACCATCTAGCGACCTCCAC-3’) in a final volume of 50 µL. The PCR cycling was performed with an initial denaturation at 98°C (30 sec), followed by 30 cycles of amplification 98°C (10 sec), 55°C (30 sec) and 72°C (1 min), and a terminal extension stage at 72°C for 10 min. PCR products were sized by electrophoresis in a 1% agarose gel containing 1× SYBR™ Safe stain (Invitrogen, ref: S33102) and 1× TAE (Tris-Acetate-EDTA) at 75V for 45 min. Amplicons of 300–1000 bp were excised from the gel and purified using E.Z.N.A. Gel Extraction Kit V-Spin (VWR). Purified amplicons were quantified by fluorescence fluorescence (Qubit™ 1× dsDNA HS Assay Kit; Thermo Fisher Scientific).

### Library preparation and genome sequencing

For TetRNAV01 virus, Illumina sequencing was performed by the Georgia Genomics Facility (now named Georgia Genomics and Bioinformatics Core) at the University of Georgia. The library was prepared using directional RNA-seq technique and sequenced using Illumina MiSeq, which produced 250 bp paired-end reads. The 3’ end of the genome was verified by using the rapid amplification of cDNA ends (3’ RACE) method. For the other four viruses (TetRNAV02–05), the double-stranded cDNA samples were sent for library preparation and sequencing at a commercial facility (SeqCenter, Pittsburgh, PA). The Illumina sequencing library was prepared using a bead-linked, transposome-based method. The library was prepared using custom 10 bp unique dual index (i7 and i5) sequences synthesized by Integrated DNA Technologies (IDT), with a target insert size of 280 bp. Multiplexed sequencing was performed on an Illumina NovaSeq X Plus sequencer producing 151 bp paired-end reads. Demultiplexing, quality control and adapter trimming was performed with bcl-convert (v4.2.4).

### Genome assembly, gene predictions and annotation

Each sample was sequenced individually using paired-end sequencing. Sequencing generated 1.83 to 2.62 million read pairs per sample, corresponding to 3.66 to 5.24 million total reads (R1 + R2). Raw reads were quality-filtered and trimmed with TrimGalore (v.6.10) (--stringency 3 --clip_R1 15 --clip_R2 15 --length 34 --quality 20) and the reads quality was evaluated with FastQC (v12.1) with default settings. Only high-quality clean reads were kept for downstream analysis. The TetRNAV01 genome was assembled from Illumina sequencing reads using SPAdes (v3.11.1) (62). For the four other viruses, independent sets of randomly down-sampled reads given a depth sequencing coverage of 100×, 500×, 1000×, and 1500× were performed using Rasusa (v2.2.2) (63). *De novo* assembly was performed using MEGAHIT (v1.2.9) (64, 65) on all sets of independently down-sampled reads and the total set of cleaned reads. The assemblies obtained with the longest contig and the smallest number of contigs were selected for subsequent analyses. The quality of single-contig viral genomes was assessed using CheckV (66). The coding sequences were predicted using getORF from EMBO (v6.6) (- minsize 600 -find 3 -find 1 - table 1 - reverse no). Functional annotations of the protein-coding genes were performed using BLASTp against the ClusteredNR database from NCBI using BLAST tool, supplemented by annotations of the conserved domains in protein sequences against CDD database using CD-search tool with setting parameters (e-value 0.01). The internal ribosome entry site (IRES) motif in the viral genome was detected using IREsite (http://www.iresite.org/) using IRES viral database (2019) (67, 68). The secondary structure and the minimum free energy (MFE) were predicted using the RNAeval webserver (v2.6.3) (69–71). The secondary structure RNA of the IGR-IRES was visualized and annotated with the tool VARNA (v3.0) (72).

### Nucleotide composition analysis

The relative frequencies of CpG and TpA dinucleotides were calculated for each viral genome using a custom Python script. First, the occurrence of each nucleotide was computed across the entire genome. Then, the observed (O) count of the CpG and TpA dinucleotides was determined by scanning the genome sequence for the exact two-nucleotide patterns, and the expected (E) dinucleotide count was based on the mononucleotide composition. Finally, the O/E ratio was calculated for each dinucleotide. A dinucleotide was considered underrepresented if its relative frequency was below 0.75.

### Genomes comparison analysis

Average nucleotide identity (ANI) between all pairs of *Tetraselmis* RNA genomes was computed using FastANI (v1.34) with a minimum fragment fraction of 0.90 (--minFraction 0.90) (73). In order to identify nucleotide polymorphic sites, the four *Tetraselmis* virus genomes (TetRNAV02, TetRNAV03, TetRNAV04 and TetRNAV05) were first aligned against the TetRNAV01 reference using MAFFT (v7.505) (74–76). Then, the multi-FASTA alignment has been implemented in the command line snp-sites tool (v2.5.1) (77) to identify single nucleotide polymorphisms (SNPs). Then, genetic variations were annotated using SnpEff (v5.4c) (78). Linear genome comparisons and polymorphism annotations were visualized using the ggplot2 package (v4.0.2) in R (v4.5.2).

### Environmental association and global distribution of RNA Tetraselmis viruses across scales

The full-length RdRP protein sequences detected in the new *Tetraselmis* viruses were blasted against the Serratus RNA viral palmprint database (79) using the palmID tool (https://serratus.io/palmid). The number of palmprint hits across environmental sources was summarized and visualized as a bar plot, while the distribution of amino acid sequence similarity across environmental sources was shown using boxplots generated with the ggplot2 (v4.0.2) package in R (v4.5.2). Latitude and longitude coordinates of the closest relative palmprint hit sequences with an identity greater than 65% were plotted on the map. A custom Python script was used to generate the map, with Pandas for data analysis and Matplotlib and Cartopy for geospatial visualization.

### Phylogenetic analysis

All phylogenetic trees were inferred using the Maximum Likelihood (ML) method implemented in IQ-TREE v2.2.2.6 (80). The best-fitting substitution model was selected using the ModelFinder and merge partitions option (-m MFP+MERGE) based on the Bayesian Information Criterion (BIC). Branch support values were estimated using 1000 replicates of both the Shimodaira-Hasegawa (SH)-like approximate likelihood ratio test (81) and the ultrafast bootstrap approximation (UFBoot) (82). Phylogenetic trees were visualized using iTOL (v7) (83).

For the near full-length 18S rRNA gene phylogeny, representative sequences of *Tetraselmis* strains were retrieved from NCBI. Three representatives of the class Trebouxiophyceae were used to root the tree. Sequences were aligned using MAFFT v7.505 (L-INS-i algorithm) (74–76) and trimmed to remove sites with ≥50% gaps using Goalign v0.3.6 (84), resulting in an alignment of 1,755 nucleotide positions.

For the RdRP domain phylogeny, closest relatives of the *Tetraselmis* viruses were identified by searching NCBI non-redundant clustered databases using the viral RdRP amino acid sequences as queries with BLASTp (e-value ≤ 1e-5). Additional RdRP sequences associated with the family *Dicistroviridae* detected in freshwater environmental samples were retrieved from Zell et al. (2022)(85). Furthermore, RdRP sequences with >70% identity previously detected in the Serratus RNA viral palmprint database (79) were added. The HMM profile of the RdRP catalytic domain (PF00680; RdRP_1) was used to extract the RdRP domain from all retrieved sequences using hmmsearch in HMMER v3.4 (--domE 1e-5). A total of 74 RdRP domain sequences were aligned with MAFFT v7.505 (L-INS-i algorithm) (74–76) and trimmed with a 50% gap cutoff using Goalign v0.3.6 (84), yielding a final alignment of 389 amino acid positions.

## Results and discussion

### Isolation, morphology, host range

Five RNA-containing viruses were isolated from surface seawater samples of Kāne‘ohe Bay, Oʻahu by challenging five closely related *Tetraselmis* sp. strains (UHM1311, UHM1313, UHM1314, UHM1315, and UHM1316) obtained from the same site. Culture UHM1316 was subsequently lost from the collection, but tests with the remaining four hosts showed that all five viruses could infect any of the other hosts, but none of the viruses could infect a somewhat more distantly related *Tetraselmis* strain (UHM1310) isolated from the same location in the bay, nor three even more divergent *Tetraselmis* strains (UHM1300, −1302, −1303) isolated from the offshore, oligotrophic waters of Station ALOHA (Fig. 1). These results indicate that the viruses have a relatively narrow host range. Electron microscopy revealed non-enveloped, untailed virions with icosahedral symmetry. The diameters of the viral particles of TetRNAV01, TetRNAV02, TetRNAV03, TetRNAV04 and TetRNAV05 were approximately 33 ± 3 nm, 34 ± 2 nm, 35 ± 2 nm, 35 ± 2 nm and 34 ± 2 nm, respectively (mean ± s.d.; n = 100 measurements per virus) (Fig. S1). These morphological features are comparable to other marine RNA viruses infecting protists previously described in the order *Picornavirales* (7, 8, 11, 13–17) (Fig. 2). The buoyant density of these viruses in iodixanol ranged from 1.138 to 1.148 g/mL (Table S2), consistent with virus densities in this medium (59).

**Fig. 1.**
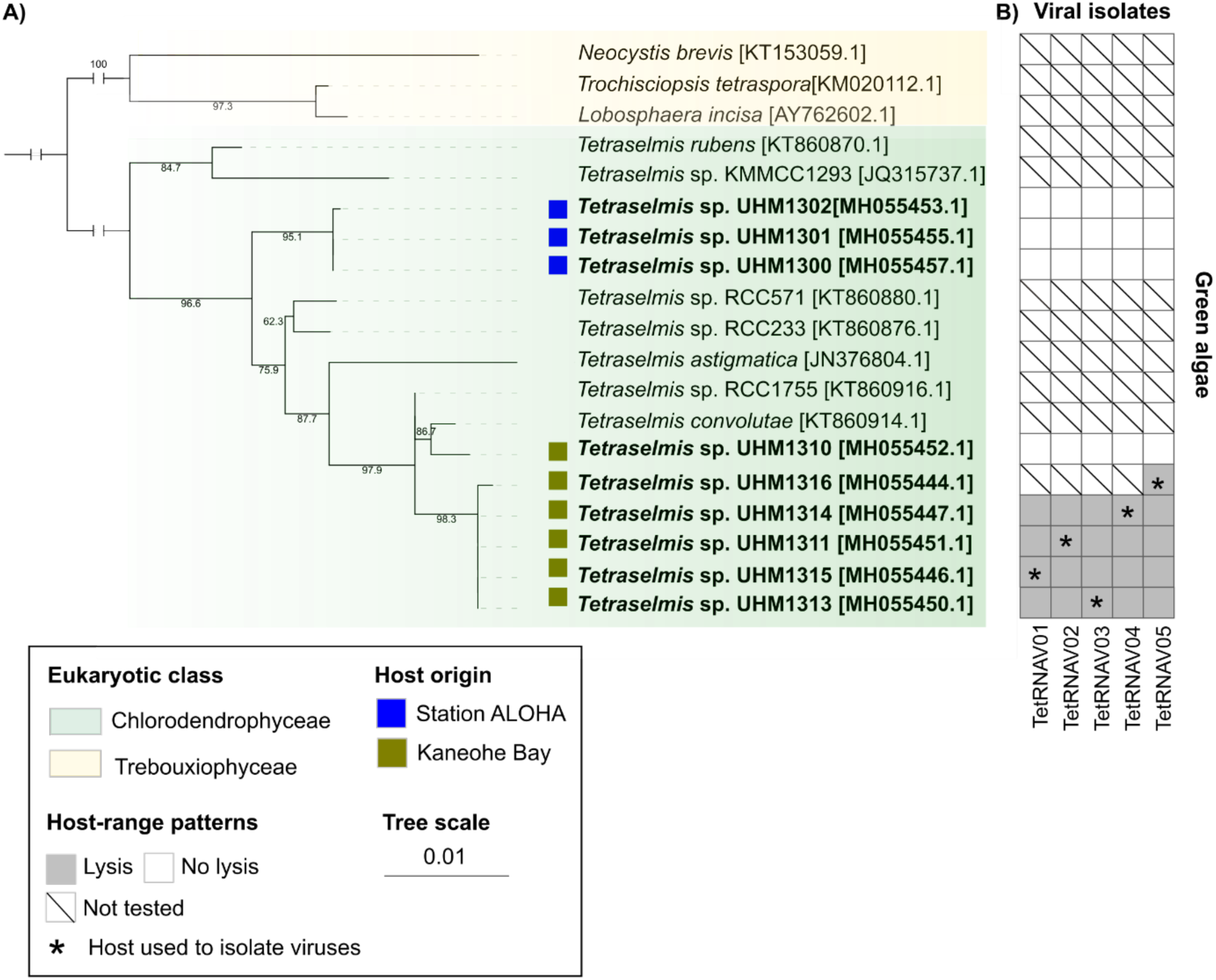
Phylogenetic relationship of phytoplanktonic strains used to assess the host range of the new isolated *Tetraselmis* viruses. A) Maximum Likelihood tree was built based on near full-length 18S rRNA gene sequences (1755 nucleotide sites) using the TN+R3 best-fit model according to Bayesian information criterion (BIC). The clade *Chlorodendrophyceae* is highlighted in green while the clade Trebouxiophyceae used as an outgroup is shown in yellow. In bold are the *Tetraselmis* strains previously isolated in Schvarcz and Steward, 2018. The geographical origin of the host strains is indicated by the square; blue from ALOHA Station (oligotrophic site) and green from Kaneohe Bay (eutrophic site). Bootstrap nodes are indicated on the branch. The tree scale bar represents the average number of substitutions per site. Branches judged to be too long in the tree were truncated and displayed with brackets for display purposes. B) The viruses tested in the cross-infection assay are shown on the horizontal axis and the *Tetraselmis* strains on the vertical axis. The cultures showing lysis by a virus are indicated by a gray filled square, an absence of lysis is indicated by an empty square. Host cultures not tested in this experiment are marked with black bars. The algal host strains used to isolate viruses are marked with a black star.

**Fig. 2.**
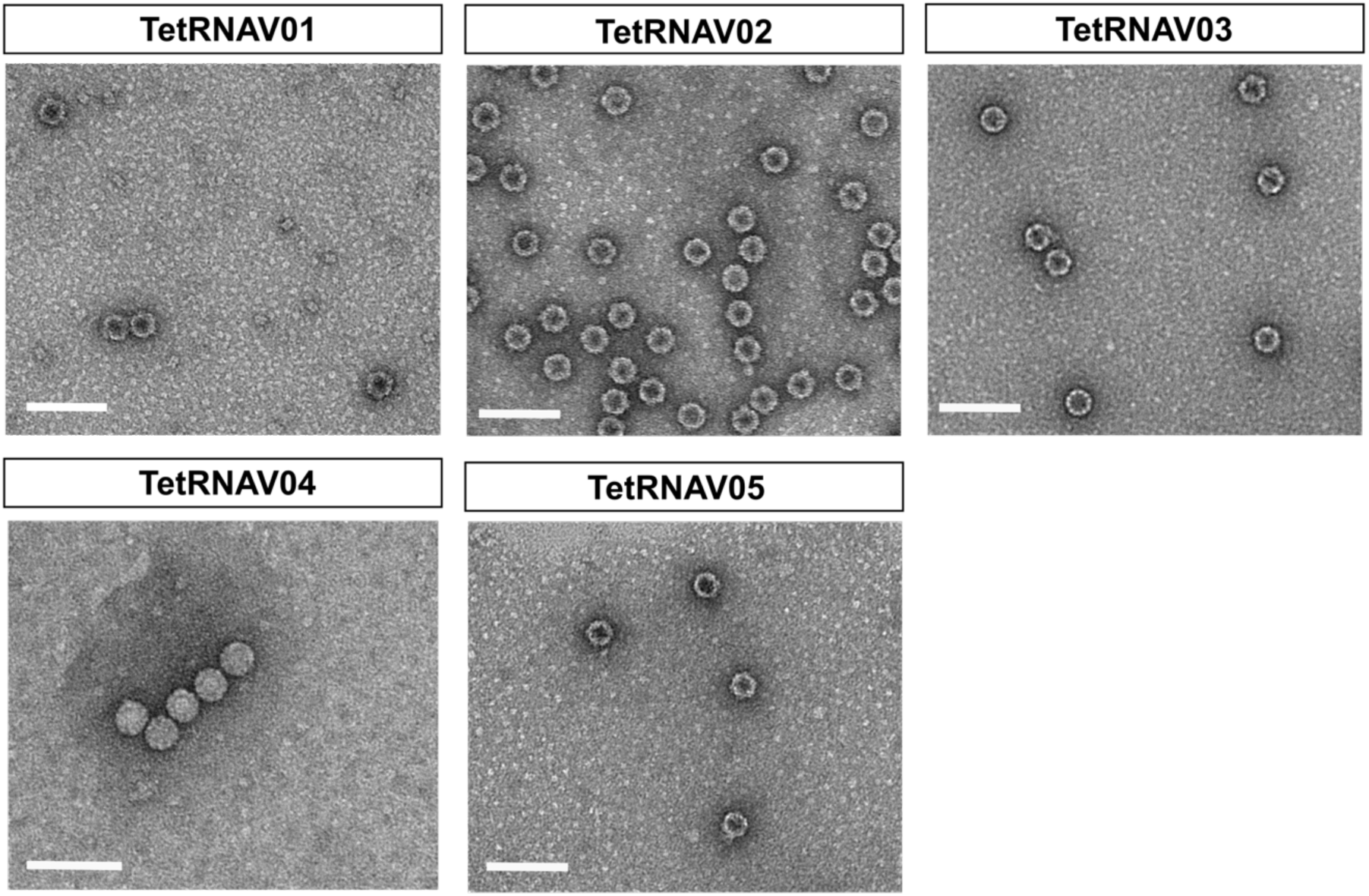
Negative staining images of the newly isolated *Tetraselmis* viral particles obtained by transmission electron microscopy. The scale bar is 100 nm.

### Genome characteristics and closest relatives

The five viruses reported in this study were initially identified as RNA-containing by their sensitivity to RNAse A, but not DNase I (Fig. S2) and were sequenced accordingly. The assembled RNA linear genome sequences of the viruses ranged in length from 9,114 to 9,225 bp (Table S2). The average base composition was estimated at 26.4% adenine, 21.45% cytosine, 22.55% guanine and 29.55% uracyl; this results in a Guanine and Cytosine (GC) content of approximately 44% (Table S2 and Table S3). The genomes consist of two non-overlapping open reading frames (ORFs), flanked by 5′- and 3′-untranslated regions (UTR) and separated by a short intergenic region (IGR), corresponding to a dicistronic genome (31, 38) (Fig. 3). ORF1 is divided into three regions encoding replication-related proteins: helicase (RNA-helicase; pfam00910) (114 aa), Tungro spherical virus-type peptidase (petidase-C3G; pfam12381) (69 aa), and RdRP (Dicistroviridae-RdRP; cd23194) (313 aa) domains. These domains form the Hel-Pro-RdRP (or Pol) core replicative module in the order *Picornavirales* (31). ORF2 encodes four structural conserved proteins which build the capsid in the order N-terminus-VP2-VP4-VP3-VP1-C-terminus (Fig. 3) (86). The VP2, VP3 and VP1 domains are the three major proteins which adopt a conserved “jelly-roll” fold important in host-virus interactions. The proteins VP2 and VP3 were homologous with a structurally conserved sequence with the capsid proteins of *Rhopalosiphum padi* virus (RhPV) (Rhv-like; cd00205; 135 aa). The protein VP4 is a small structural protein located at the N-terminal end of the VP3 protein, which is cleaved from the precursor VP0 to produce the mature protein in the families *Dicistroviridae* and *Iflaviridae* (87–89). The VP4 shared sequence homology with the Dicistro-VP4 domain (46 aa; PF11492) and VP1 with a C-terminal CRPV capsid domain (CRPV-capsid; PF08762; 207 aa), both related with capsid proteins in *Cricket paralysis* virus (family *Dicistroviridae*). The BLAST searches revealed that both proteins were closely related to the Beihai picorna-like virus 79 (BHPLV-79) (Genbank accession: YP_009333536) with an identity of amino acid sequences of approximately 42%. This virus was originally sequenced from penaeid shrimp tissue and was phylogenetically associated to the family *Dicistroviridae* (order *Picornavirales*) (33).

**Fig. 3.**
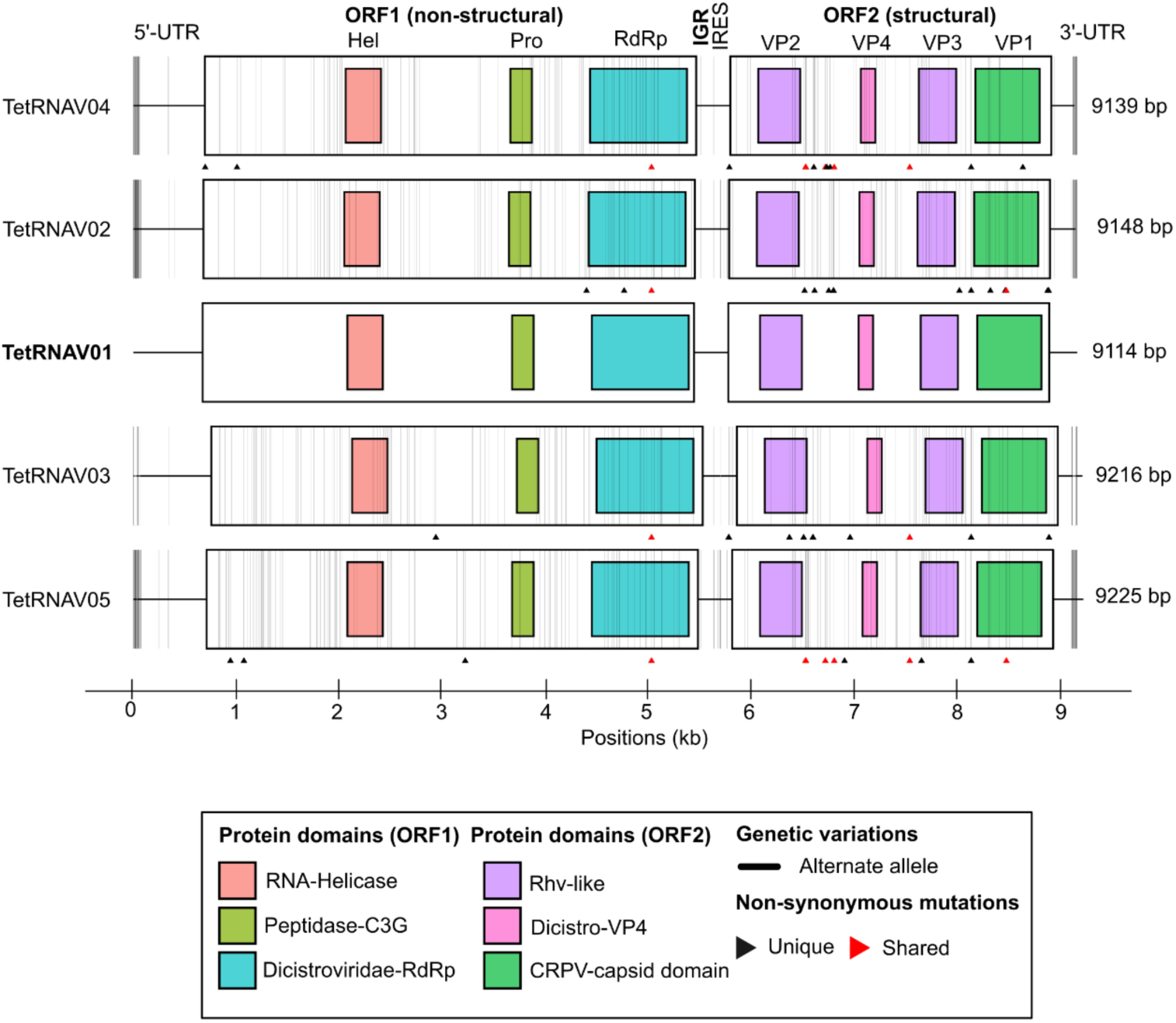
Comparative linear genome map of the *Tetraselmis* virus isolates. The genomes are organized from longest (bottom) to shortest (top). 5′-UTR and 3′-UTR denote the 5′ and 3′ untranslated regions, respectively. *Tetraselmis* viruses consist of two open reading frames (ORFs) which both translate polyproteins and are separated by a small intergenic region (IGR). ORF1 encodes a non-structural protein, the RNA-dependent RNA polymerase (RdRP), whereas ORF2 encodes the structural capsid protein. Functional domains within each ORF are shown as colored boxes: RNA-helicase (red), peptidase-C3G (green), Dicistroviridae-RdRP domain (blue), RhV-like superfamily (purple), Dicistro-VP4 (pink), and CRPV-capsid (dark green). An internal ribosome entry site (IRES) was detected in the IGR of each genome. Vertical black bars indicate alternate alleles relative to the reference genome (TetRNAV01). Arrows display positions of non-synonymous mutations; mutations unique to a single genome are shown in black, whereas those shared by at least two genomes are shown in red. Genome coordinates are indicated in kilobases (kb).

### Replication mechanisms and host-virus associations

A computational analysis revealed a short internal ribosome entry site (IRES) element (58 nt) located within the IGR, whose nucleotide sequence is conserved across all TetRNAV genomes (Fig. 3, Fig. S3A). The best match corresponded to the IGR-IRES of *Rhopalosiphum padi virus* isolate M1 (RhPV; KX610810), an aphid-infecting virus member of the genus *Cripavirus*, with 88% nucleotide sequence identity (E-value 4e-09) (90). The matching sequence is homologous to the prominent stem-loop structure III and IV characterized in the RhPV IGR-IRES (91, 92) and very similar to the IV and V stem-loop segments in the secondary structure predicted in the *Plautia stali* intestine virus (93). Three short, conserved RNA segments were identified in the IGR-IRES sequence. These conserved motifs are located at similar positions in the stem-loop segments of the secondary structure in other invertebrate picornavirads (93) (Fig. S3B). It has been proposed that these factors play a key role in IRES activity and could represent the site of ribosome entry in these viruses (91, 93). In dicistrovirids, translation is typically mediated by two IRES elements upstream of the initiation codon: one in the 5′-UTR upstream of ORF1, which directs the synthesis of nonstructural proteins, and another within the IGR, which drives translation of the capsid protein (37, 39). The IGR-IRES can imitate both a translation elongation factor and the anticodon loop of a tRNA paired with mRNA, enabling translation to occur without the need for canonical initiation factors. This unique mechanism represents an alternative strategy for ribosome recruitment to mRNA, establishing a distinct mechanism of translation initiation, allowing them to control the translation of each ORF independently (39). In contrast, only the IGR-associated IRES was identified in the TetRNAV genomes, and its sequence is considerably shorter than those reported in other dicistroviruses (e.g., ∼58 nt compared to ∼200 nt). The predicted secondary structure of this IGR-IRES, with a minimum free energy of ΔG ≈ −12.5 kcal/mol, consists of two distinct stem–loop domains separated by a short, unpaired linker (Fig. S3C). The 5′ domain forms a compact hairpin with internal bulges, whereas the 3′ domain is more complex, comprising an extended stem with internal loops and a deeply nested helical region (Fig. S3C). Notably, no pseudoknot structures were detected in the TetRNAV IRES, despite their essential role in ribosome recruitment and stabilization during the initiation of translation in dicistroviruses (94). Although two conserved stem-loop structures were found associated with three conserved RNA segments, the TetRNAV IRES differs substantially from those described in other dicistroviruses (37, 39, 94–99). Furthermore, the presence of a canonical AUG (Methionine – Met) start codon at the beginning of the capsid ORF suggests that translation initiation likely involves initiator Met-tRNA, in contrast to dicistrovirus IGR-IRES elements that initiate from non-AUG codons *via* pseudoknot-mediated ribosome positioning (91, 93). Together, the absence of canonical pseudoknot features and the use of AUG start codon could indicate a divergent mechanism of translation initiation that likely does not rely on direct ribosome mimicry. Instead, these observations support a non-canonical, potentially novel atypical factor-dependent IRES mechanism regulating structural protein synthesis as observed in the Bemisia-Associated Dicistrovirus 2 (100). Finally, the mechanism underlying RdRP (nonstructural protein) translation in TetRNAV genomes remains unclear. The IRES in picornavirads and dicistrovirids appears to be diverse, consisting of different class variants that involve various mechanisms for initiating translation (97, 101, 102). Unusual IRES architectures have been reported in metagenome-assembled genomes of marine dicistrovirids (102–104), which suggests that this divergence may be an adaptation to specific hosts or environmental conditions.

Finally, in order to gain insight into the genomic signatures of the TetRNAVs and virus-host associations, we calculated the frequencies of dinucleotides across the genomes. The CpG (Avg ratio = 0.66) and TpA (Avg ratio = 0.74) observed/expected (O/E) ratios indicated only moderate underrepresentation (suppression) of CpG and TpA dinucleotides (Fig. S4). Dinucleotide suppression was generally observed to be stronger than this in vertebrate- and plant-infecting viruses (105, 106). The patterns observed in TetRNAV genomes are consistent with previous reported studies in invertebrates viruses (107). This pattern may reflect an adaptation to the host cellular replication machinery, shaped by complex evolutionary pressures, including immune evasion systems, as observed across RNA viruses (106).

### Intraspecific comparison of Tetraselmis RNA virus genomes

Pairwise ANI values among the five viral genomes ranged from 97.5% to 98.3%, supporting their classification as distinct strains within the same species (Table S4) (108). Using TetRNAV01 as the reference genome, we identified a total of 453 single nucleotide polymorphisms (SNPs), corresponding to approximately 5% of nucleotide positions across the genome (Fig. 3). For the substitutions located in the coding regions, the majority were synonymous (n = 302; 66%), whereas 42 were non-synonymous. Notably, 34 (80%) of the non-synonymous mutations were unique to a single genome (Fig. 3), likely suggesting an independent strain-level genome diversification. A cluster of non-synonymous mutations was identified in the VP2 C-terminal sequence in the viral genomes, one of the major structural (capsid) viral proteins in the *Picornavirales* (31). This domain is therefore likely to be important for binding to the host receptor, making it subject to host-virus coevolution. Finally, the remaining variants (n = 109) were located in intergenic regions and therefore did not affect protein-coding sequences.

A base substitution analysis revealed a high transition-to-transversion ratio (Ts/Tv) of 8.65, indicating that nucleotide substitutions were strongly biased toward transitions. The very low dN/dS ratios observed for the virus-encoded RNA-dependent RNA polymerase (RdRP; dN/dS = 0.046) and capsid proteins (dN/dS = 0.30) provide evidence of strong purifying (negative) selection, reflecting the functional constraints acting on these essential viral proteins. Such selection pressure on both structural and non-structural domains was also recently described in marine metagenome-assembled genomes of viruses classified within the family *Manaviridae* (109). These results support the suggestion that RdRP domain is under purifying selection that maintains the functionality of the active sites of the protein (110).

### Evolutionary relationships within the Dicistroviridae

A phylogenetic reconstruction based on the RdRP domain and capsid protein placed the newly isolated TetRNAVs within the family *Dicistroviridae*, order *Picornavirales* (Fig. 4a and 4b). Consistent with the RdRP and capsid phylogenies, the concatenated protein phylogeny recovered the same topology (Fig. S5), indicating a conserved evolutionary signal across the genome and supporting a congruent evolutionary history of the TetRNAVs. At present, there are three defined genera in this family: *Aparavirus*, *Cripavirus*, and *Triatovirus* (99). Acute bee paralysis virus (ABPV) is the exemplar virus of the genus *Aparavirus* (111) and Cricket paralysis virus (CrPV) is the exemplar virus of the genus *Cripavirus* (112). *Triatoma virus* (TrV), which was initially classified as a member of the *Cripavirus* genus (113), has since been reassigned as the founding member of the genus *Triatovirus* (114). All of the characterized viruses within these genera infect arthropods (39). However, sequences derived from metatranscriptomes of aquatic animal tissues and metagenomes targeting RNA viruses in aquatic samples have revealed additional diverse sequences that cluster with these genera. The hosts for these environmental sequences are unknown.

**Fig. 4.**
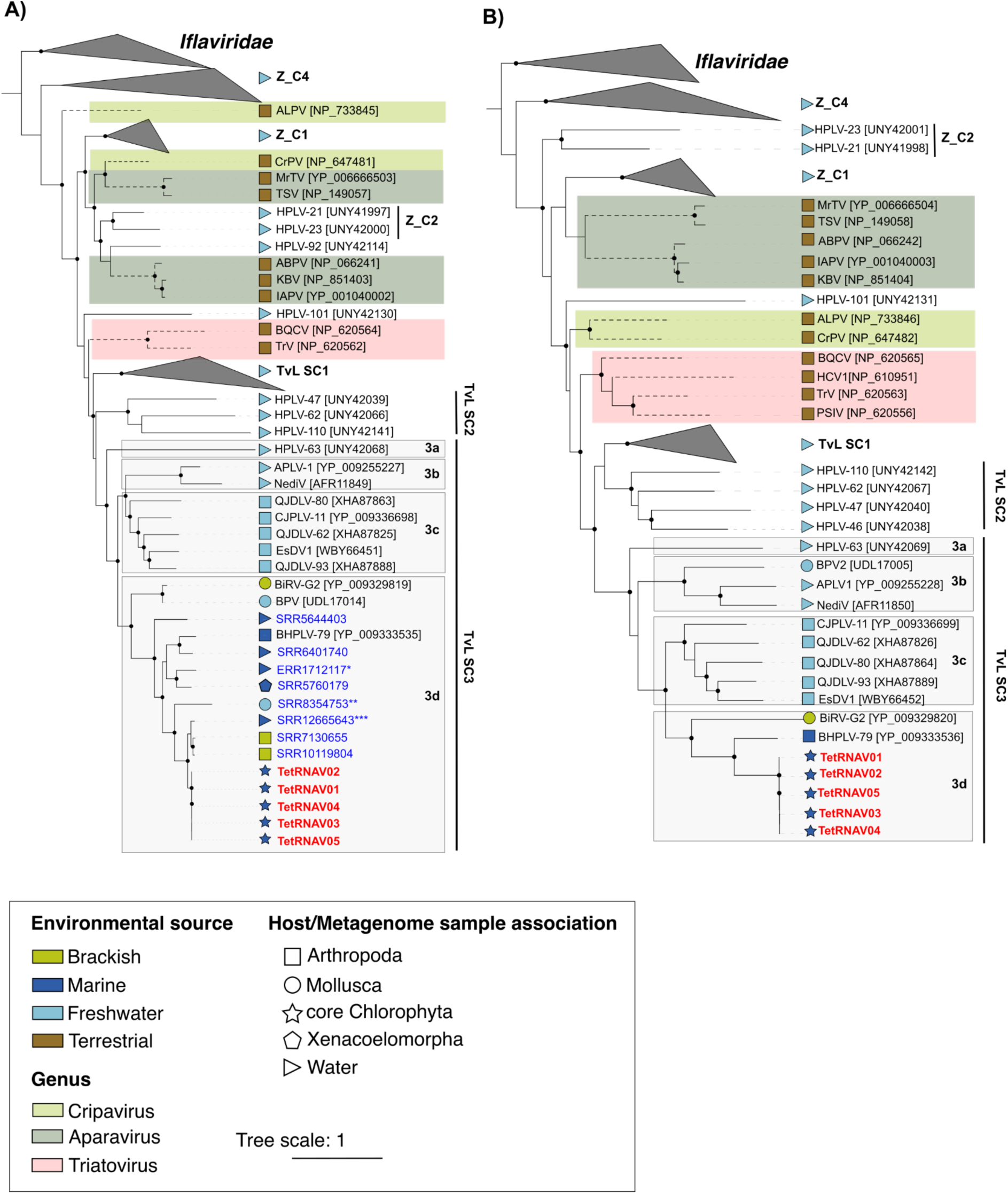
Evolutionary relationships of the newly isolated *Tetraselmis* viruses within the family *Dicistroviridae*. A) A maximum-likelihood phylogenetic tree was inferred from 380 amino acid positions of the RNA-dependent RNA polymerase (RdRP) domain and B) from 750 amino acid positions of the full-length capsid protein alignments. Members of the family *Iflaviridae* were used as an outgroup to root the tree. Representative sequences from isolated viruses within the genera *Cripavirus*, *Aparavirus*, and *Triatovirus* are indicated by dotted branches. The sample association from which each viral sequence was obtained is indicated by symbols: squares indicate Arthropoda, circles Mollusca, ellipses core Chlorophyta, icosahedra Xenacoelomorpha, and triangles environmental water samples. The environmental source is indicated by color: green and dark blue represent marine environments, light blue freshwater, and brown terrestrial environments. The genus *Cripavirus* is shown in light green, *Aparavirus* in dark green, and *Triatovirus* in red. The closest environmental-hit sequences identified in public datasets from the Serratus RNA viral palmprint database are highlighted in blue, whereas the viruses isolated in this study are shown in bold red. The best-fit substitution models (A; Q.PFAM+F+I+R5 and B; Q.PFAM+F+R5) were selected according to the Bayesian Information Criterion (BIC). Nodes with bootstrap support over 75% are shown as filled circles in the tree. The scale bar represents the average number of substitutions per site. Virus abbreviations and their associated protein accession numbers are listed in Table S6a. Identical Serratus RNA viral palmprint amino acid sequences clustering on the same branch are indicated as follows: *SRR9108850; **SRR8354754 and SRR8354751; ***SRR12661218 and SRR12661217. TvL SCx: Triatovirus-like virus subclade. Z_Cx: Clades containing environmental sequences derived from freshwater in Germany that were previously assigned to dicistrovirus-like clusters in Zell et al. (85).

The TetRNAV sequences, together with environmental sequences from marine and freshwater habitats, formed a broad but well-supported sister clade to the officially classified triatoviruses, referred to as the *Triatovirus*-like virus (TvL) large subclade (Fig. 4). The TvL was divided into three major lineages, TvL-SC1, TvL-SC2, and TvL-SC3, which were separated by well-supported internal branches. TvL-SC1 and TvL-SC2 comprised several sequences previously recovered from lake water samples in Germany and designated Havel picorna-like viruses (HPLVs) (85). Both lineages occupied basal positions within the TvL phylogeny, suggesting an early divergence from the common ancestor of the group. In contrast, TvL-SC3 represented a more diverse lineage comprising sequences associated with freshwater, marine, and brackish environments.

TvL-SC3 was further divided into four well-supported subclades (3a–3d), highlighting substantial evolutionary diversification within this lineage. TvL-SC3a–c comprised previously characterized HPLV-related viruses together with environmental sequences and were predominantly associated with freshwater environments. These lineages included a diverse assemblage of viruses associated with aquatic arthropods, such as Qianjiang dicistro-like virus (QJDLV)-93, QJDLV-80, QJDLV-62, and Changjiang picorna-like virus 11 (CJPLV-11), which were recovered from tissues of the red swamp crayfish Procambarus clarkii collected in freshwater environments in China (33). The group also included *Eriocheir sinensis* dicistrovirus 1 (EsDV1), identified in freshwater crabs affected by mitten disease. Antarctica picorna-like virus 1 (APLV-1), detected in Lake Limnopolar, Antarctica, was reported as one of the most abundant RNA viruses in this polar aquatic ecosystem (115), whereas the closely related Nedicistrovirus (NediV) was recovered from untreated sewage water in Nepal (116). Together, these viruses formed a well-supported basal lineage within TvL-SC3, further supporting the presence of an early-diverging, predominantly freshwater-associated group.

In contrast, the TetRNAVs clustered within the more derived sister subclade TvL-SC3d. Whereas TvL-SC3a–c was dominated by freshwater-derived sequences, TvL-SC3d was predominantly composed of viruses associated with marine environments. This lineage included BHPLV-79, originally identified from penaeid shrimp tissues in China (33). Within this marine-associated clade, *Biomphalaria pfeifferi*-associated virus (BPV), identified from RNA extracted from homogenized whole-snail *Biomphalaria pfeifferi* tissue (117), and the closely related bivalve-associated sequence BiRV-G2 (118) formed an early-diverging lineage, while the TetRNAVs occupied a more derived position within SC3d.

### Ecological, Evolutionary, and Geographical Patterns of Related Viral Lineages

We investigated whether RNA virus sequences infecting *Tetraselmis* sp. could be detected in publicly available sequencing datasets that had not been specifically analyzed for viruses. To do so, we performed BLAST searches of the RdRP protein sequences against the Serratus RNA viral palmprint database using the palmID tool. This search identified a total of 480 RdRP palmprint hits with protein sequence identities ranging from 53% to 89% (Table S5). These sequences originated from different environmental sources and geographically distant regions worldwide (Fig. 5). Most of the best matches were recovered from aquatic environments, and the majority of those were derived from seawater samples (n = 84) (Fig. 5A). The highest palmprint identities (89% amino acid identity) were detected in crab hepatopancreas tissues from *Helice tientsinensis* (SRR7130655) and *Scylla paramamosain* (SRR10119804), both collected in China (Guangzhou and Yancheng, respectively) (Fig. 5B). Three other homologs with 89% amino acid identity were identified in seawater samples collected at Marineland in Florida (USA). All three of the latter sequences originated from the same metagenomic dataset but corresponded to different incubation conditions (SRR12665643; SW_R4_TP1: incubated in the dark at 28 °C for 29 hours, SRR12661218; SW_R1_TP0: initial timepoint of incubation, SRR12661217; SWP_R3_TP0: initial timepoint of incubation).

**Fig. 5.**
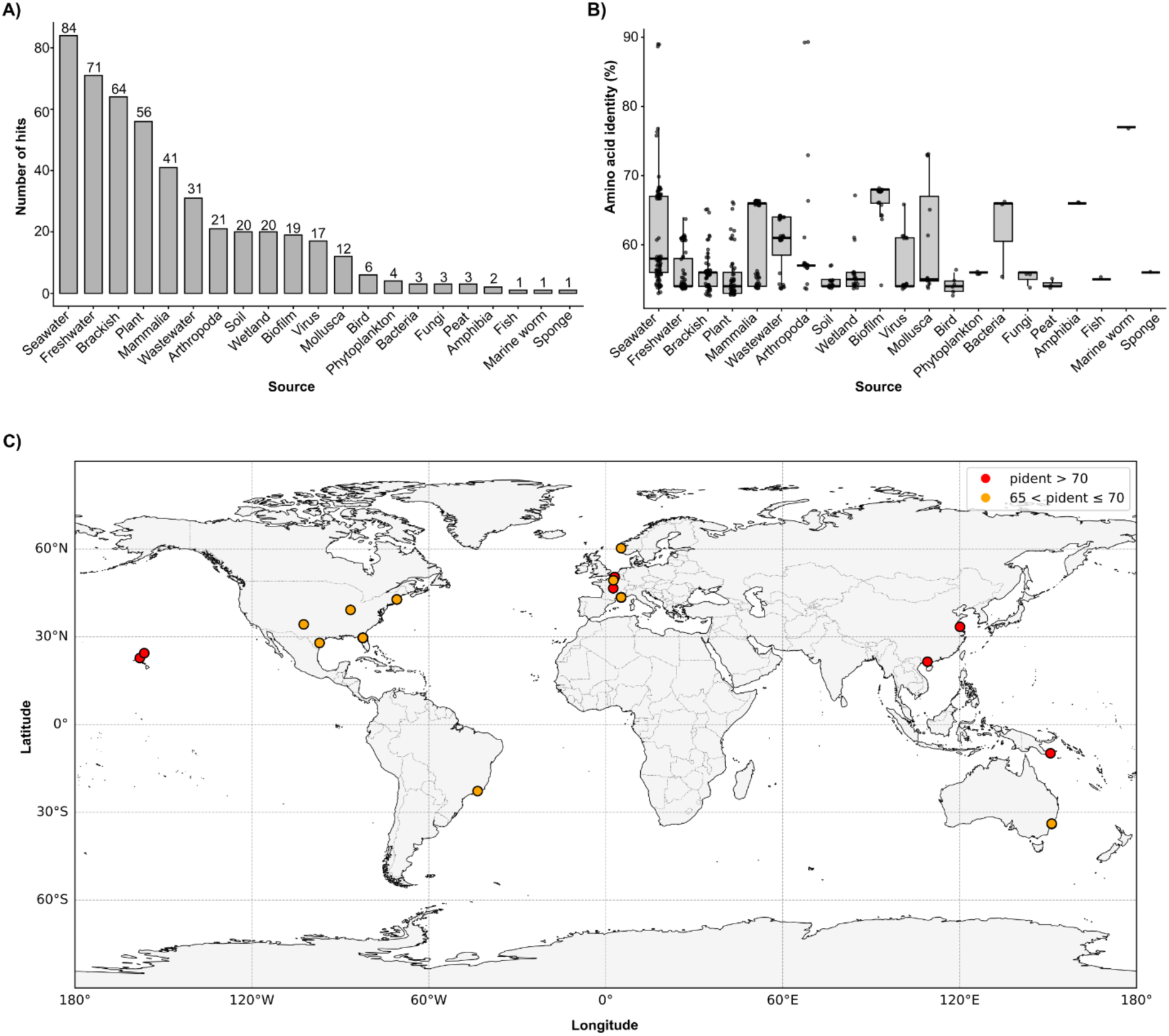
Geographic distribution of related *Tetraselmis* viral lineages based on RdRP palmprint sequences. A) Bar plot showing the number of palmprint hits detected in environmental datasets that are related to the RdRP sequences of the isolated viruses. Bars are ordered by the total number of hits. B) Boxplot illustrating the distribution of amino acid sequence similarity between environmental palmprint hits and the RdRP sequences of the isolated viruses. C) Global geographic distribution of environmental samples from which related palmprint sequences were identified. Sequences showing 65–70% amino acid identity are indicated in yellow, whereas sequences with >70% identity are shown in red. Raw data for the palmprint hits—including SRA accession, palmID, amino acid sequence identity, and environmental source information—are available in the supplemental data (Table S5).

Two additional closely related viral lineages (76% amino acid identity) were detected in the North Pacific Subtropical Gyre near Station ALOHA, only a few kilometers offshore from where the new *Tetraselmis* RNA viruses were isolated. The first of these gyre palmprint sequences was identified in seawater samples collected and size-fractionated between 180 and 20 µm during the *Tara Oceans* expedition in 2011 at station TARA_131 (SRA accession: ERR1712117), as part of a meta-transcriptomic survey targeting unicellular eukaryotes. The second closely related RdRP palmprint sequence was also recovered from a seawater sample (sample name: S21C1_02) collected in 2015. The latter sample was prefiltered through 100 µm Nitex mesh and collected onto a 0.2 µm pore-size polycarbonate filter from the surface mixed layer (∼15 m depth). This sample (SRA accession: SRR9108850), as part of Diel1 eukaryotic metatranscriptome project, was collected during R/V Kilo Moana cruise KM1513 (SCOPE HOE-Legacy 2) in July 2015 (119).

Also intriguing is a palmprint hit sharing 77% sequence identity was detected in the sequenced genome dataset of the acoel flatworm *Symsagittifera roscoffensis* isolated in France (SRA accession: SRR5760179) (120). This marine flatworm-like animal, belonging to the phylum Acoela, lives in an obligate symbiotic relationship with the alga *Tetraselmis convolutae* during its adult stage (47, 121). Notably, the *Tetraselmis* sp. strains used to isolate the RNA viruses in this study share 99.3% 18S rRNA sequence identity with *Tetraselmis convolutae* strain RCC1564 (isolated in Roscoff (France) from the Atlantic Ocean) and branch together in a strongly supported clade (Fig. 1). Together, these results indicate that RNA viruses infecting *Tetraselmis* and related to the *Dicistroviridae* may be widespread in marine environments. The detection of a related sequence in a dataset from *Symsagittifera roscoffensis* raises the possibility that such viruses may occur within the *S. roscoffensis* holobiont, potentially through their association with its algal symbiont *T. convolutae*. The phylogenetic reconstruction based on the RdRP domain placed all of these sequences from aquatic samples and the isolated TetRNAV genomes in a diverse cluster with a branch point that suggests a common ancestor with triatoviruses (Fig. 4a). Notably, the SRR7130655 and SRR10119804, are the closest relative homologues to the TetRNAVs (89% of amino acids identity), and are both derived from mud crab tissues.

Finally, four palmprint hits, sharing 56% sequence identity, were detected in datasets associated with the ciliated protist *Stentor polymorphus* (family *Stentoridae*, order *Heterotrichida*). This organism, which lives in a symbiotic association with green algae (specifically *Chlorella*) (122, 123), was found in sediment samples collected from a pond water in Uppsala (Sweden; 59°50′19.0″N, 17°37′21.4″E) with an aim to assess transcriptional changes during cell regeneration (124). The sequences were detected under four experimental conditions: posterior bisection at 300 min post-cell split (SRR6714495), posterior bisection at 90 min post-cell split (SRR6714489), and two non-regenerating control samples (SRR6714476 and SRR6714475). Several symbiotic associations have been well documented, including the relationship between *Chlorella variabilis* and the mixotrophic ciliate *Paramecium bursaria* (125), within which a giant virus infecting *Chlorella* (*Paramecium bursaria* Chlorella virus 1 — PBCV-1) was originally discovered (126). To date, no RNA virus infecting *Chlorella* has been reported. Both *Tetraselmis* and *Chlorella* belong to the core chlorophytes within the division Chlorophyta (40, 41); however, only DNA viruses—either double- or single-stranded—have so far been described in *Tetraselmis* (52, 53, 127–129). In this study, we report the first RNA viruses isolated from the green alga *Tetraselmis*. Furthermore, the detection of an environmental sequence closely related to TetRNAV in the ciliated protist *Stentor polymorphus* in our analysis suggests the existence of environmental RNA viral lineages potentially capable of infecting *Chlorella* within the holobiont. These data highlight that *Dicistroviridae*-related RNA viruses infecting protists are widely distributed, showing high host specificity and likely spanning a broad range of phytoplankton taxa.

## Conclusion

In this study, we report the isolation and characterization of five closely related, but distinct, positive-sense, single-stranded RNA (+ssRNA) viruses that infect a narrow lineage of green algae within the *Tetraselmis* genus. These appear to be the first ssRNA virus isolates infecting members of the phylum Chlorophyta, but a search of RNA-dependent RNA polymerase (RdRP) sequences in publicly available environmental datasets reveals other TetRNAV-like lineages widely distributed across aquatic ecosystems. The presence of an atypical IGR-IRES in the genomes suggests that TetRNAVs use a non-canonical translation initiation mechanism, potentially reflecting adaptation to specific host factors or a strategy that promotes viral protein synthesis at the expense of host translation. The virion morphology— tailless, non-enveloped capsids approximately 35 nm in diameter—was typical of other protist-infecting RNA viruses, but we were surprised by the phylogenetic analyses that placed these protist-infecting viruses within the family *Dicistroviridae*, all the classified members of which infect arthropods. Considering the clustering of TetRNAV protein sequences with many others derived from aquatic environments to form a well-supported monophyletic group adjacent to the *Triatovirus* genus, these new isolates may serve as the basis for defining a new genus of alga-infecting viruses within the *Dicistroviridae*. The detection of related sequences in material derived from diverse aquatic animal tissues —marine arthropods and mollusks—and also from water samples, suggests that some inferred virus-animal connections may reflect indirect links arising from virus-infected protists associated with the animal. Our results expand the breadth of known alga-infecting RNA viruses and illustrate the value of continued isolation and characterization of marine virus-host systems for interpreting environmental sequence data.

## Data availability

The five *Tetraselmis* RNA virus genomes were deposited in Genbank under accessions numbers: TetRNAV01, PZ360873; TetRNAV02, PZ360874; TetRNAV03, PZ360875; TetRNAV04; PZ360876, TetRNAV05, PZ360877. Bioproject: PRJNA1458800

## Acknowledgements

This work was supported by NSF awards OCE-2129697 and RII Track-2 FEC 1736030 (to G.F.S. and K.F.E.), NSF award OCN 0826650 (to G.F.S), Simons Foundation award 566853 (to K.F.E), and EF 0424599 (to G.F.S.). We thank Dr. Orion S. Rivers at the University of Hawaiʻi at Mānoa Biological Electron Microscope Facility for his assistance and guidance with electron microscopy procedures. We also thank the personnel in the Hawai’i Ocean Time-series program for assistance with water collection. The technical support and advanced computing resources from the University of Hawaii Information Technology Services – Cyberinfrastructure, funded in part by the National Science Foundation CC* awards # 2201428 and # 2232862, are gratefully acknowledged.

## Author contributions

J.T., K.F.E. and G.F.S. conceptualized the project; K.F.E. and G.F.S. acquired funding; J.T., K.F.E., G.F.S., C.R.S. and K.A.M. conducted the investigations; J.T., K.F.E., G.F.S. and C.R.S. provided resources and supervised the project; J.T., K.F.E. and G.F.S. wrote the paper with input from all authors.

## Ethics declarations

The authors declare no competing interests.

## Supplemental data

Table S1. Summary of the phytoplankton *Tetraselmis* spp. strains challenged for isolation of viruses from environmental seawater. **EC enrichment cultures of whole or filtered seawater samples amended with f/2 media. SD, serial dilution of the algal strain, performed using three consecutive rounds of serial dilution.

**Table S2.** Summary of the phenotypic and genomic features of TetRNAV viruses.

| Virus name | Capsid size (nm) (n = 100) | Buoyant density Iodixanol (g/mL) | Assembled contigs | Genome size (bp) | GC content (%) | #ORFs |
| --- | --- | --- | --- | --- | --- | --- |
| TetRNAV02 | 34 ± 2 | 1.138 | 1 | 9,148 | 44.1 | 2 |
| TetRNAV03 | 35 ± 2 | 1.148 | 1 | 9,216 | 43.9 | 2 |
| TetRNAV04 | 35 ± 2 | 1.138 | 1 | 9,139 | 43.9 | 2 |
| TetRNAV05 | 34 ± 2 | 1.142 | 1 | 9,225 | 44.2 | 2 |
| TetRNAV01* | 33 ± 3 | 1.135 | 1 | 9,114 | 43.8 | 2 |

**Table S3.** Nucleotide composition of the *Tetraselmis* RNA virus genomes. This was calculated by counting the occurrence of each individual nucleotide (A,T,G and C) in the genome sequence and dividing by the total number of nucleotides and multiplied by hundred to express as a percentage. The average nucleotide composition across all genomes is reported (average). A: adenine; C: cytosine, G: guanine, T: tyrosine.

|  | TetRNAV05 | TetRNAV04 | TetRNAV03 | TetRNAV02 | TetRNAV01 | Average |
| --- | --- | --- | --- | --- | --- | --- |
| %A | 26.32 | 26.51 | 26.5 | 26.42 | 26.49 | 26.448 |
| %C | 21.59 | 21.41 | 21.46 | 21.49 | 21.3 | 21.45 |
| %G | 22.66 | 22.52 | 22.45 | 22.56 | 22.58 | 22.554 |
| %T | 29.43 | 29.55 | 29.59 | 29.53 | 29.63 | 29.546 |

**Table S4.** Pairwise of average nucleotide identity (ANI) of *Tetraselmis* RNA virus genomes computed with FastANI (Jain et al., 2018). The values are expressed as a percentage.

|  | TetRNAV03 | TetRNAV04 | TetRNAV02 | TetRNAV05 | TetRNAV01* |
| --- | --- | --- | --- | --- | --- |
| TetRNAV03 | 100 |  |  |  |  |
| TetRNAV04 | 98.03 | 100 |  |  |  |
| TetRNAV02 | 97.62 | 97.86 | 100 |  |  |
| TetRNAV05 | 97.70 | 97.92 | 97.61 | 100 |  |
| TetRNAV01* | 97.72 | 98.26 | 97.88 | 97.50 | 100 |

Table S5. List of conserved sub-sequences within viral RdRP (palmprint) hits homologous to the novel *Tetraselmis* viral lineages, identified through Serratus palmID analysis against RdRP palmprint database (PALMdb). Information includes Sequence Read Archive (SRA) accession, percentage of amino acid identity (pident), PalmID barcodes, origin, and origin categories (Category).

Table S6. a) List of viruses used to build a single-gene phylogenetic tree based on the RdRP protein sequence. b) List of viruses used to build multi-gene concatenated phylogenetic trees based on the both RdRP and capsid protein sequences These tables provide information on the virus’s origin (sample/organism) and/or host (host), host taxonomy (phylum; subphylum/class), and environment (source) in which it was collected.

---

**Fig. S1.**
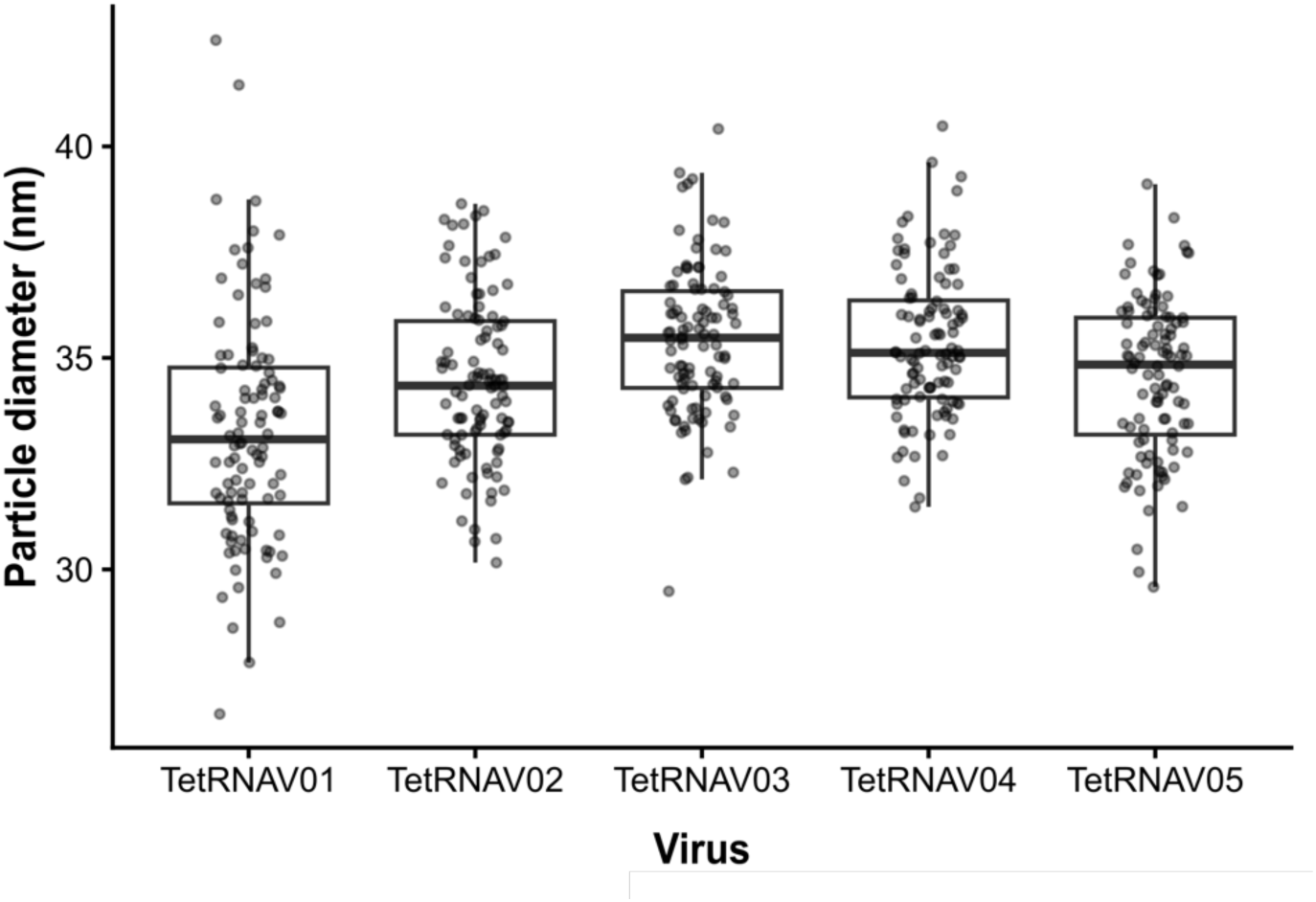
Box plots represent the distribution of particle diameters measured by transmission electron microscopy (n = 100 particles per isolate). Boxes indicate the interquartile range, horizontal lines represent the median, and points show individual measurements.

**Fig. S2.**
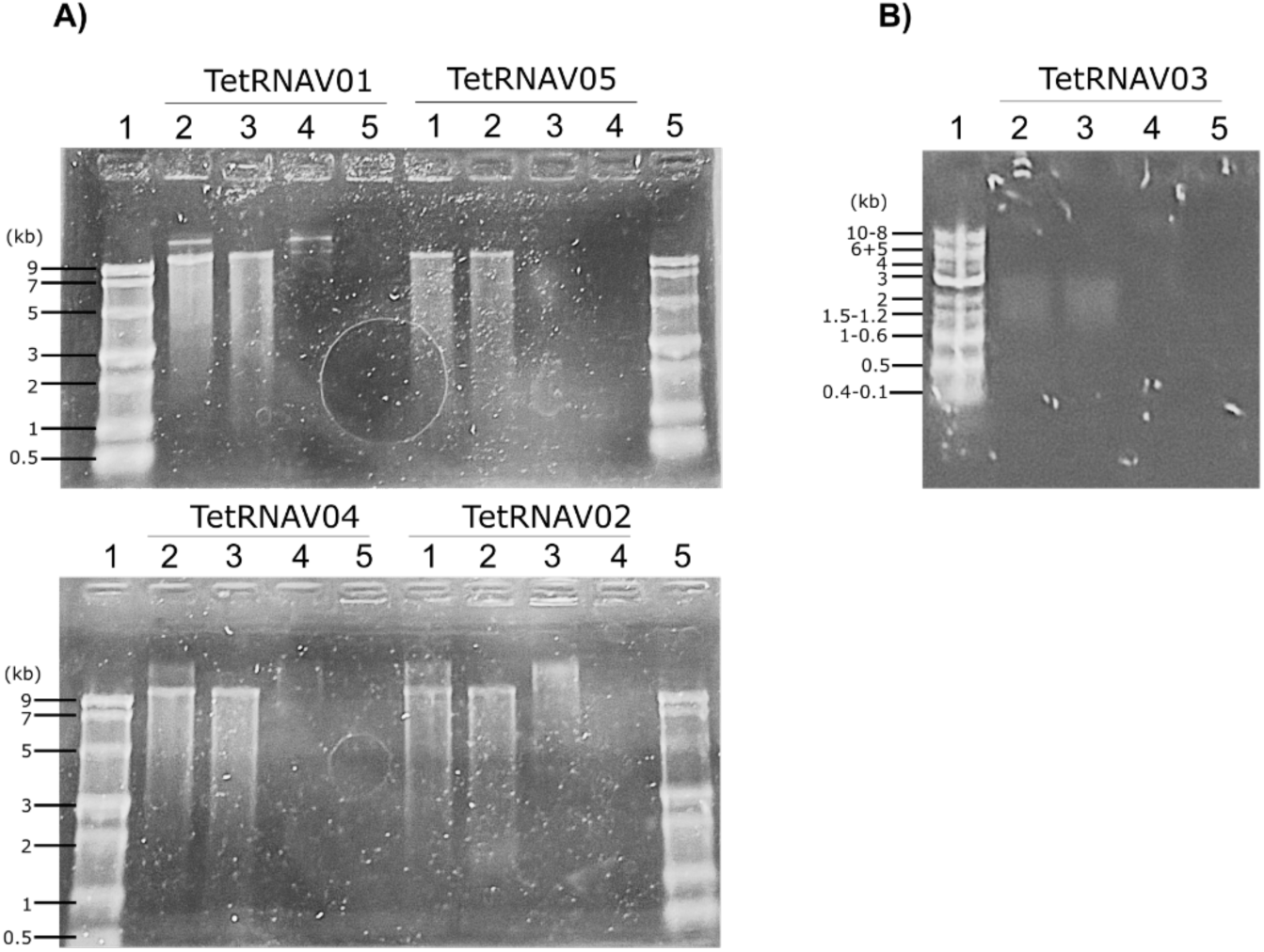
Electrophoresis gel of endonuclease-digested and undigested TetRNAVs genomic RNA. A) TetRNAV01, TetRNAV02, TetRNAV04 and TetRNAV05 genomic RNA was digested with DNAse I and/or RNAse and loaded onto a 1% agarose-formaldehyde gel. Lane 1: ssRNA Ladder. Lane 2: control (Genomic RNA was not subjected to enzymatic digestion). Lane 3: DNAse I digestion. Lane 4: RNAse A digestion. Lane 5: DNAse I and RNAse A digestion. B) TetRNAV03 genomic RNA was digested with DNAse I and/or RNAse and loaded onto a 0.8% agarose gel. Lane 1: Quick-Load® Purple 1 kb Plus DNA Ladder. Lane 2: control (Genomic RNA was not subjected to enzymatic digestion). Lane 3: DNAse I digestion. Lane 4: RNAse A digestion. Lane 5: DNAse I and RNAse A digestion. kb: kilobases. Note that in Panel A, samples were run on a denaturing gel with an RNA ladder. In the samples for TetRNAV01, −02, and −05, the largest band likely represents residual, unresolved host genomic DNA contamination. The band just above the top 9 kb marker is intact viral genomic RNA, and the smear below degraded RNA. In Panel B, samples were run on a non-denaturing gel with a DNA ladder, and the lower bands from 1–3 kb were presumed to be the viral genomic RNA not properly running according to size because of secondary structure.

**Fig. S3.**
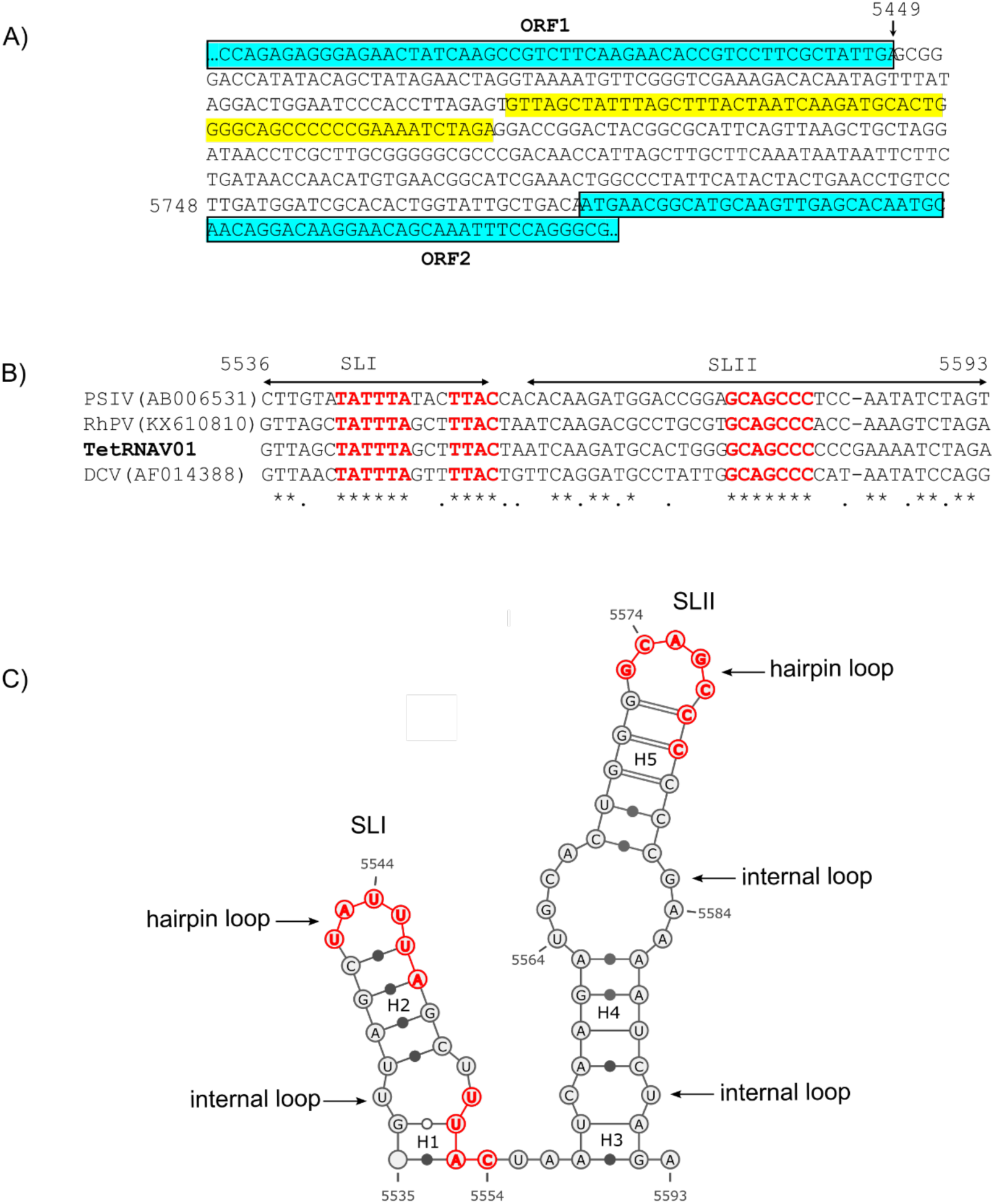
Characterization of the IGR-IRES in the TetRNAV genome. (A) Partial nucleotide sequence showing the IRES region (yellow) located within the intergenic region (IGR) between the two ORFs (blue). A multiple sequence alignment of the region upstream of the capsid-coding ORF2 is presented, using TetRNAV01 as the reference (bold), alongside dicistroviruses *Rhopalosiphum padi* virus (RhPV; KX610810), *Plautia stali* intestine virus (PSIV; AB006531), and *Drosophila* C virus (DCV; AF014388). Asterisks (*) indicate nucleotides conserved across all four viruses, while dots (.) indicate partial conservation. Conserved short sequence motifs, thought to play important roles in IRES activity, are highlighted in red. Predicted stem–loop (SLx) elements in the TetRNAV01 secondary structure are indicated by double-headed arrows above the sequence. (C) Line drawing of the predicted secondary structure of the TetRNAV01 RNA containing the IRES domain. Conserved short RNA segments shared among the four viruses are highlighted in red, and nucleotide positions are indicated. Each helix predicted within the steem-loops was numbered (Hx). The structure was generated using Visualization Applet for RNA (VARNA)(v3-93) (https://varna.lisn.upsaclay.fr/).

**Fig. S4.**
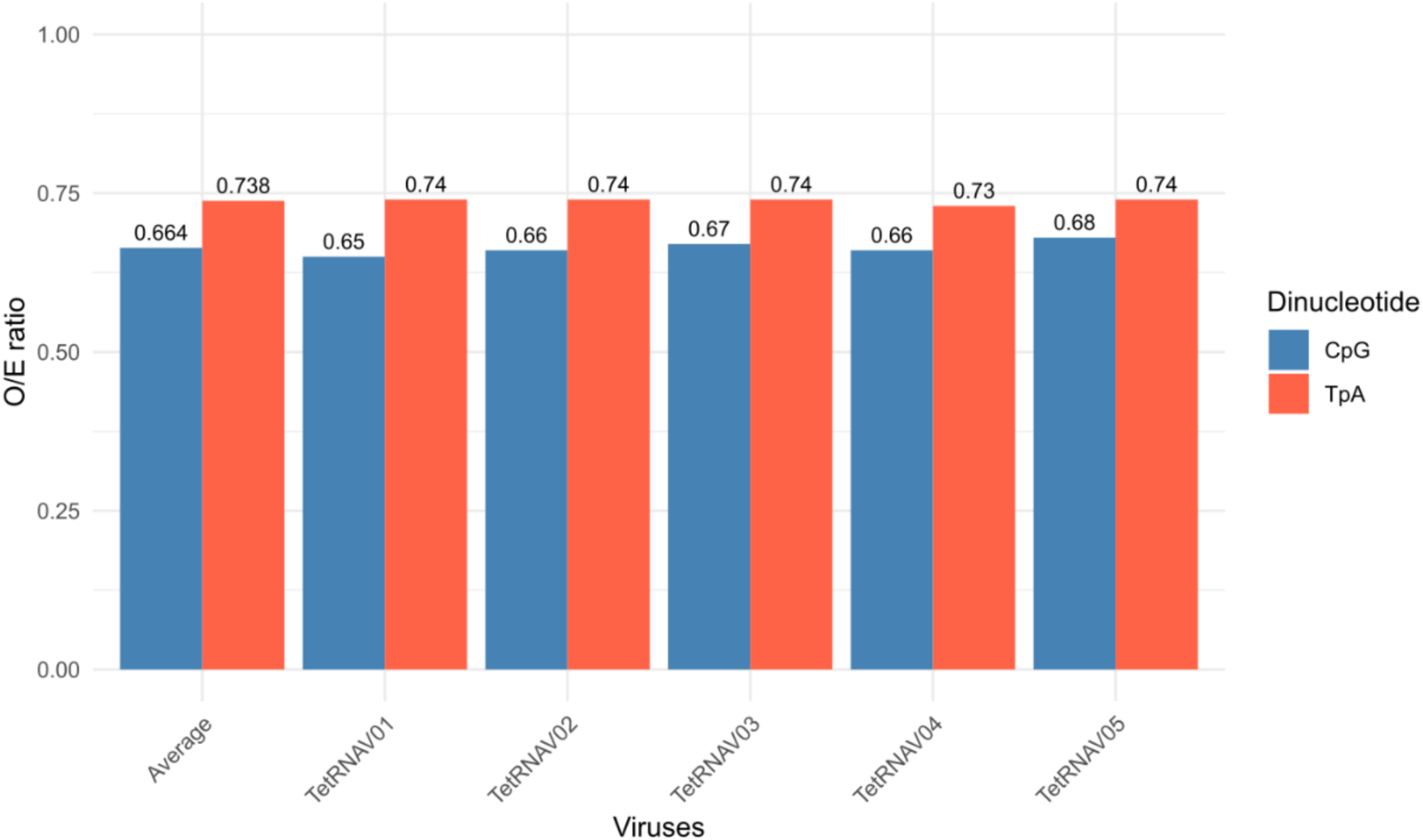
Dinucleotide frequencies of CpG and TpA in *Tetraselmis* RNA virus genomes. The observed number (O) corresponds to the actual count of a dinucleotide in the sequence, while the expected number (E) is derived from the frequencies of individual nucleotides under the assumption of random association. Observed-to-expected (O/E) dinucleotide ratios were calculated using the number of CpG and TpA occurrences across all overlapping positions in each sequence (L−1, where L is the sequence length), divided by their expected frequencies estimated from nucleotide composition as (C × G)/L for CpG and (T × A)/L for TpA, with C, G, T, and A representing the counts of each nucleotide. An O/E ratio < 0.75 indicates CpG underrepresentation (or suppression) and selection against TpA.

**Fig. S5.**
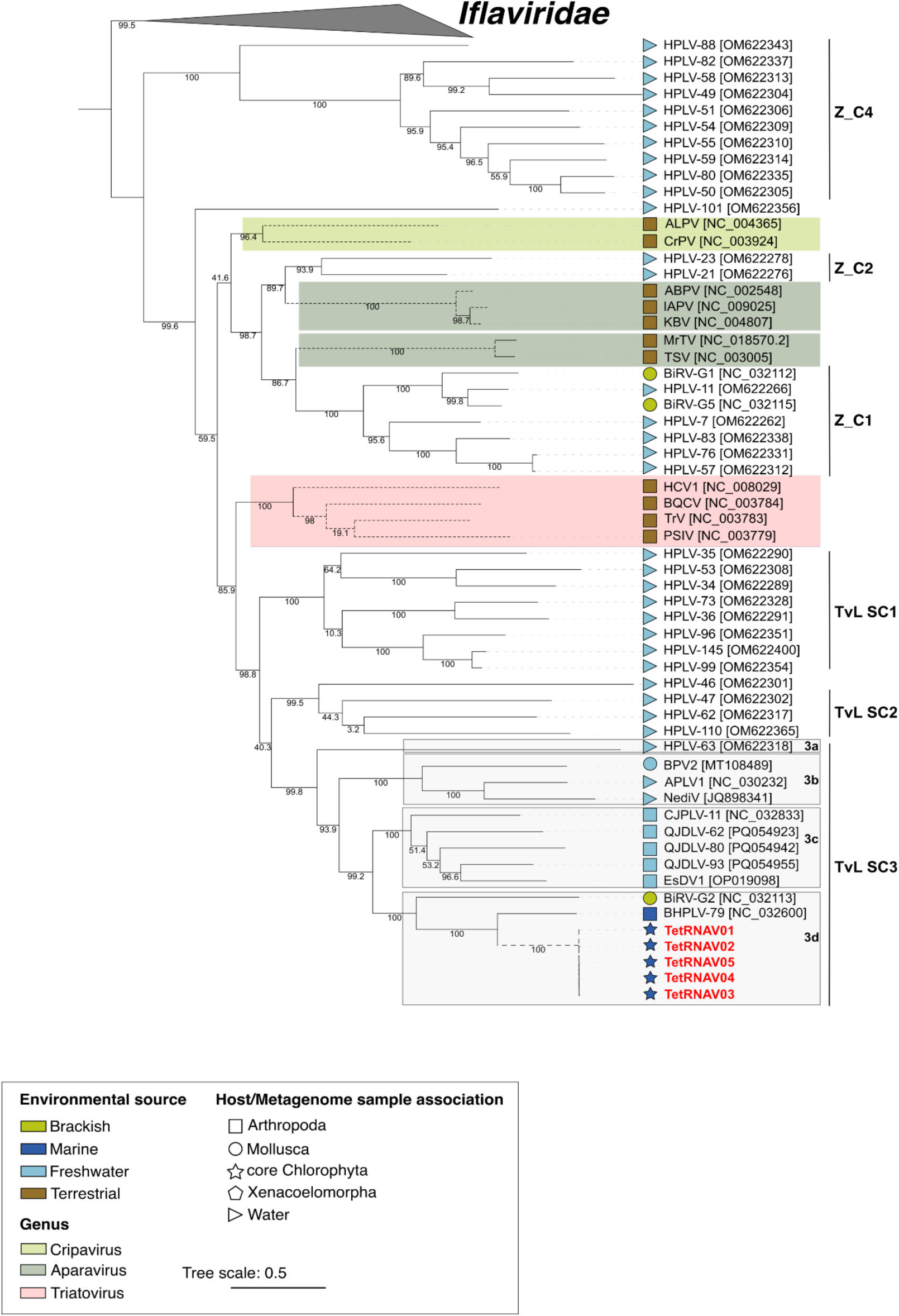
Maximum-likelihood phylogenies inferred from a concatenated amino acid alignment of the complete capsid protein and the RdRP domain of the non-structural polyprotein (1140 amino acid positions). The best-fit substitution models selected according to the Bayesian Information Criterion (BIC) were Q.PFAM+F+I+R5. Members of the *Iflaviridae* family, including the Deformed Wing Virus (DWV), Sacbrood Virus (SBV), Infectious Flacherie

