## Supplementary figures and images for "First Evidence of Dicistroviruses Infecting Protists"

### Supplemental Figure 1

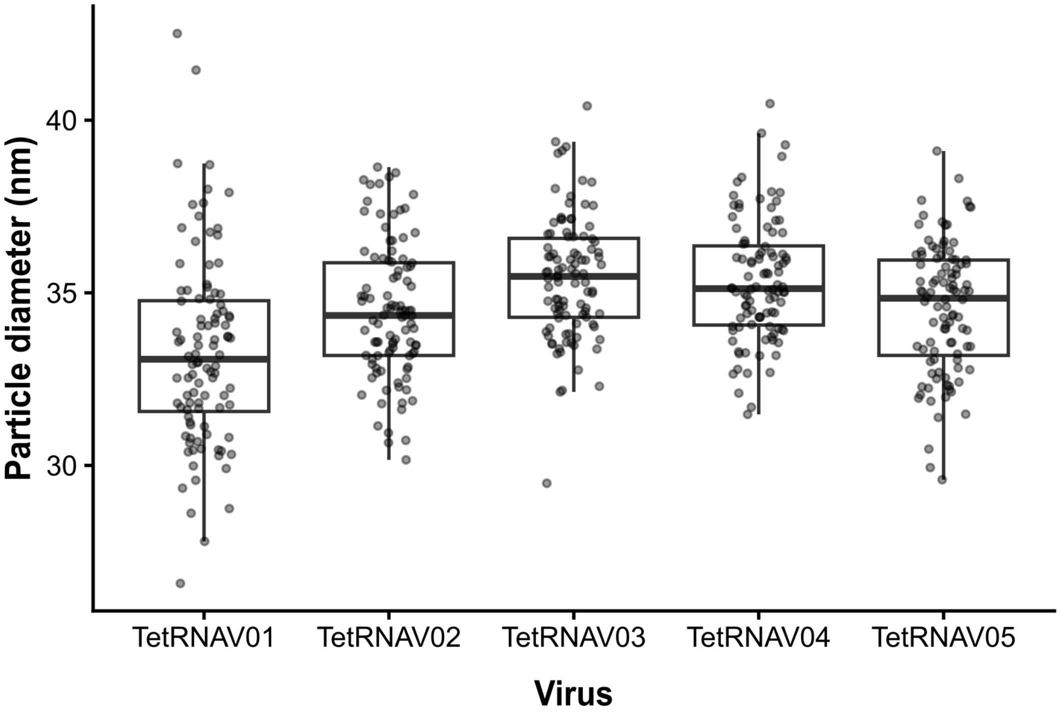

### Supplemental Figure 2

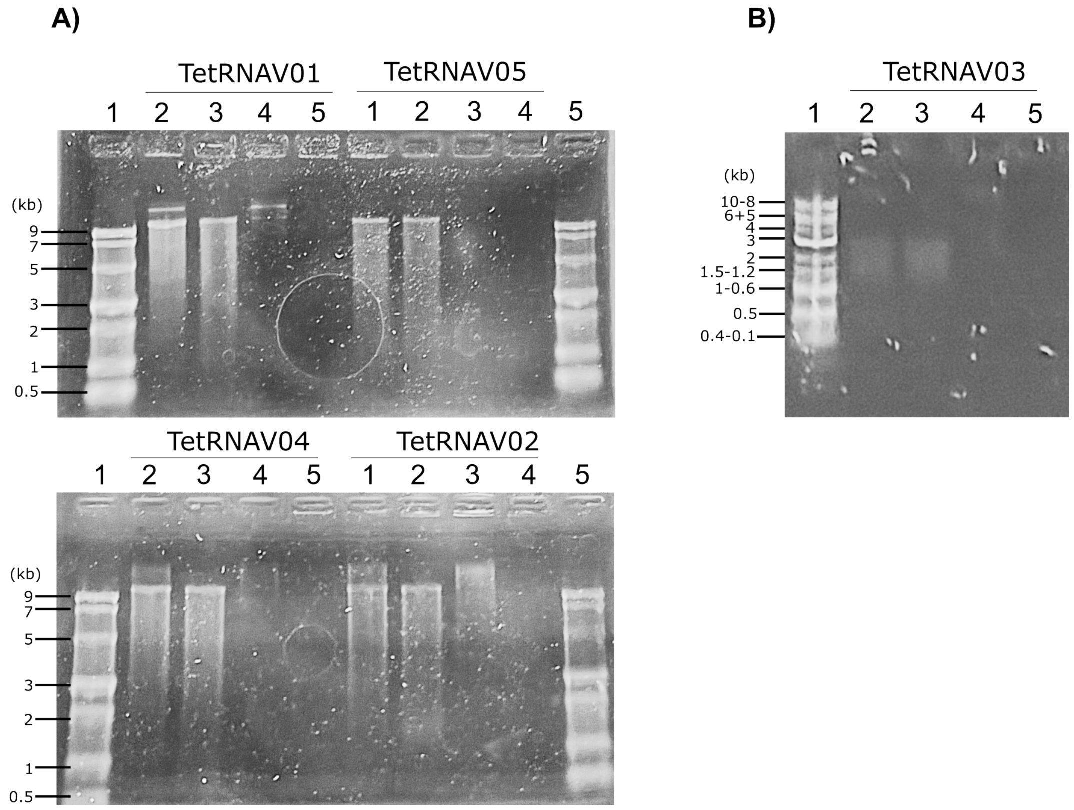

### Supplemental Figure 4

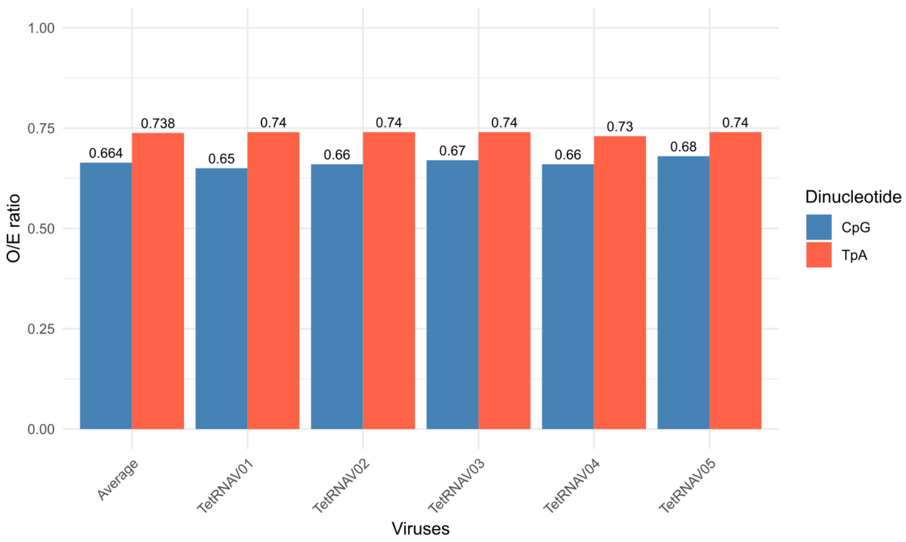

### Supplemental Figure 5

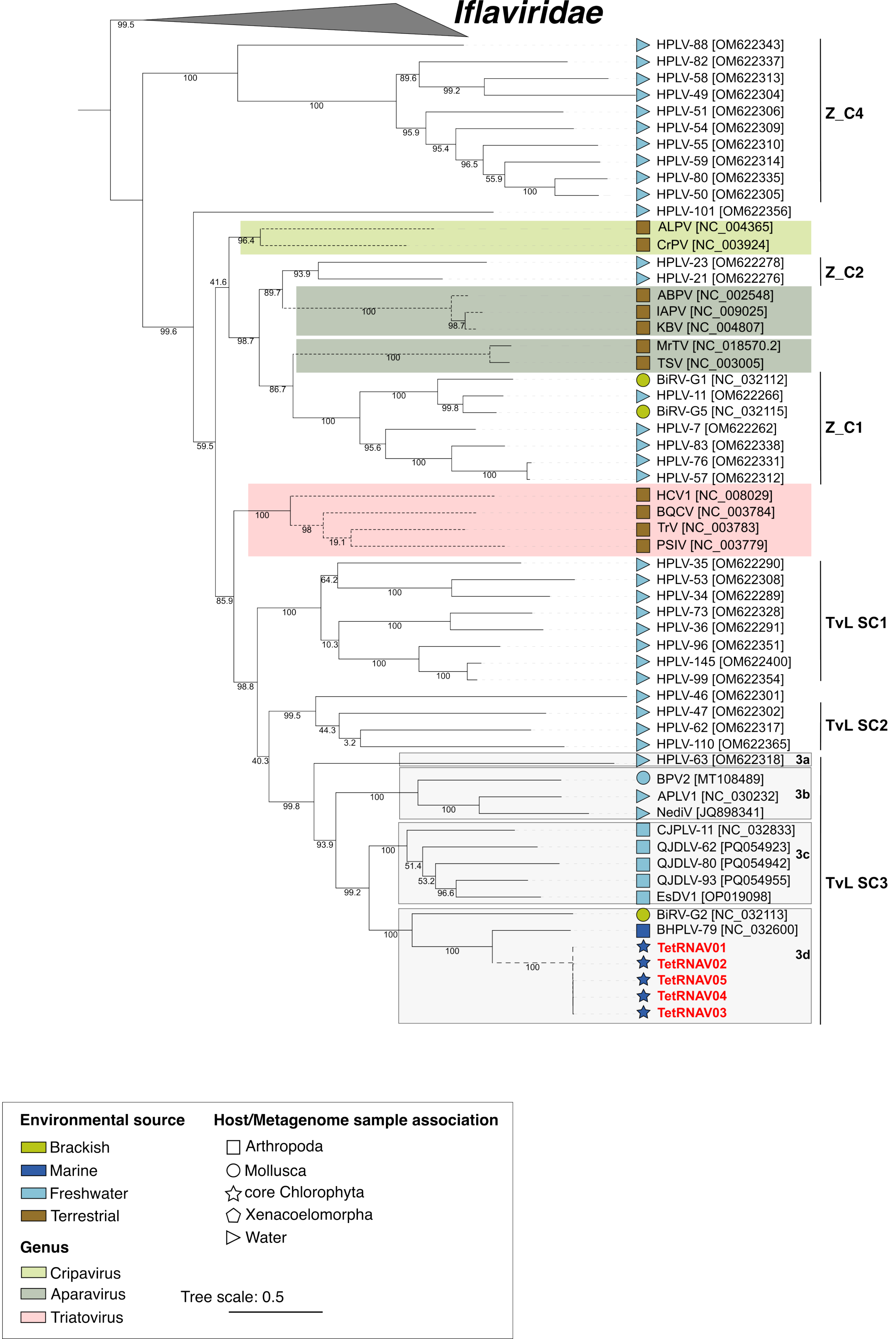
