## Supplemental Figure 3 for "First Evidence of Dicistroviruses Infecting Protists"

A)

```
                                                                    5449
                                     ORF1                              ↓
..CCAGAGAGGGAGAACTATCAAGCCGTCTTCAAGAACACCGTCCTTCGCTATTGAGCGG
GACCATATACAGCTATAGAACTAGGTAAAATGTTCGGGTCGAAAGACACAATAGTTTAT
AGGACTGGAATCCCACCTTAGAGTGTTAGCTATTTAGCTTTACTAATCAAGATGCACTG
GGGCAGCCCCCCGAAAATCTAGAGGACCGGACTACGGCGCATTCAGTTAAGCTGCTAGG
ATAACCTCGCTTGCGGGGGCGCCCGACAACCATTAGCTTGCTTCAAATAATAATTCTTC
TGATAACCAACATGTGAACGGCATCGAAACTGGCCCTATTCATACTACTGAACCTGTCC
5748 TTGATGGATCGCACACTGGTATTGCTGACAATGAACGGCATGCAAGTTGAGCACAATGC
AACAGGACAAGGAACAGCAAATTTCCAGGGCG..
                 ORF2
```

B)

```
                5536       SLI                              SLII                        5593
PSIV(AB006531) CTTGTATATTTATACTTACCACACAAGATGGACCGGAGCAGCCCTCC-AATATCTAGT
RhPV(KX610810) GTTAGCTATTTAGCTTTACTAATCAAGACGCCTGCGTGCAGCCCACC-AAAGTCTAGA
TetRNAV01      GTTAGCTATTTAGCTTTACTAATCAAGATGCACTGGGGCAGCCCCCCGAAAATCTAGA
DCV(AF014388)  GTTAACTATTTAGTTTTACTGTTCAGGATGCCTATTGGCAGCCCCAT-AATATCCAGG
                **.   ******   .****..  **.**.*   .     *******   . **  .**.**
```

C)

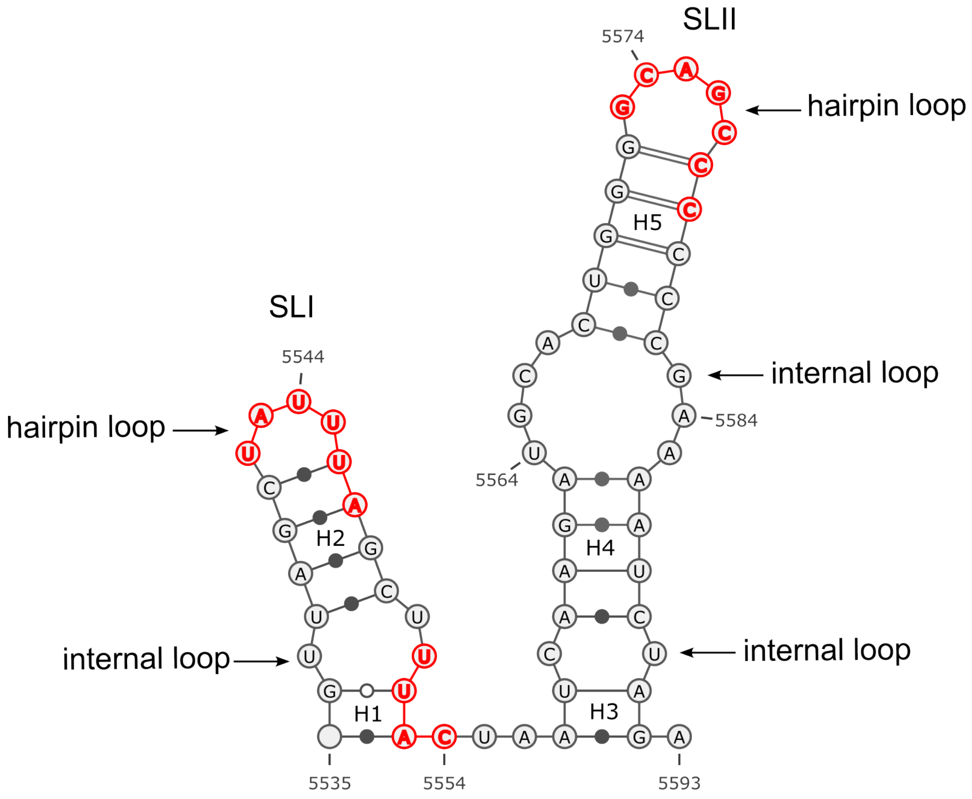
